# Non-Invasive Quality Control of Organoid Cultures Using Mesofluidic CSTR Bioreactors and High-Content Imaging

**DOI:** 10.1101/2024.07.19.604365

**Authors:** Seleipiri Charles, Emily Jackson-Holmes, Gongchen Sun, Ying Zhou, Benjamin Siciliano, Weibo Niu, Haejun Han, Arina Nikitina, Melissa L. Kemp, Zhexing Wen, Hang Lu

**Affiliations:** Interdisciplinary Program in Bioengineering, Georgia Institute of Technology, 311 Ferst Drive NW, Atlanta, Georgia 30332, U.S.A.; Wallace H. Coulter Department of Biomedical Engineering, Georgia Institute of Technology, 313 Ferst Drive NW, Atlanta, Georgia 30332, U.S.A.; School of Chemical & Biomolecular Engineering, Georgia Institute of Technology, 311 Ferst Drive NW Atlanta, Georgia 30332, U.S.A.; Departments of Psychiatry and Behavioral Sciences, Cell Biology, and Neurology, Emory University School of Medicine, 615 Michael Street, Atlanta, Georgia 30322, U.S.A.; Graduate Program in Molecular and Systems Pharmacology, Laney Graduate School, Emory University, 615 Michael Street, Atlanta, GA, 30322, U.S.A.; School of Biological Sciences, Georgia Institute of Technology, 310 Ferst Drive NW, Atlanta, Georgia 30332, U.S.A.

**Keywords:** brain organoids, microfluidics, bioreactors, label-free imaging, machine learning assisted quality control, 3D culture, growth dynamics

## Abstract

Human brain organoids produce anatomically relevant cellular structures and recapitulate key aspects of *in vivo* brain function, which holds great potential to model neurological diseases and screen therapeutics. However, the long growth time of 3D systems complicates the culturing of brain organoids and results in heterogeneity across samples hampering their applications. We developed an integrated platform to enable robust and long-term culturing of 3D brain organoids. We designed a mesofluidic bioreactor device based on a reaction-diffusion scaling theory, which achieves robust media exchange for sufficient nutrient delivery in long-term culture. We integrated this device with longitudinal tracking and machine learning-based classification tools to enable non-invasive quality control of live organoids. This integrated platform allows for sample pre-selection for downstream molecular analysis. Transcriptome analyses of organoids revealed that our mesofluidic bioreactor promoted organoid development while reducing cell death. Our platform thus offers a generalizable tool to establish reproducible culture standards for 3D cellular systems for a variety of applications beyond brain organoids.

## 1. Introduction

The lack of robust and efficient human-specific models that resemble the human brain has long been a significant challenge for studying human brain development, modeling brain diseases, and testing drug efficacy. With the rapid development of stem cell technology, 3D brain organoids can be generated to recapitulate critical organ and tissue-specific features of cell assembly, integration, and organization in the developing brain and exhibit human-relevant properties not observed in animal models.^[1–7]^ They provide a unique opportunity to study human organ development, model brain diseases, and screen therapeutics.^[2–4, 8–14]^ Despite the exciting potential of brain organoids, their large-scale applications have been limited due to several critical challenges. First, 3D brain organoids need a long-term culture (weeks to months) to develop human-relevant tissue structures and can grow from a few hundred microns to millimeters. The long culturing time and large range of organoid sizes make it especially difficult to maintain efficient oxygen and nutrient exchange while minimizing cell-damaging shear stress by using bulk-scale bioreactors (e.g., spinner flasks, orbital shakers).^[2, 15]^ Second, organoids vary widely in the diversity of cell types present and structural features in culture.^[16]^ To date, the variability can only be assessed by end-point assays. Although culturing protocols to improve uniformity have been proposed, the lack of automated and non-invasive assessment of live organoids still complicates quantitative analyses and limits the applicability of brain organoids in disease modeling and drug screening applications.^[17–19]^ Various microfluidic technologies have been developed to improve the consistency and quality of organoid culture. In recent years, these technologies have been used to reduce laborious manual manipulation and the risk of external contamination.^[20–29]^ The reduced footprint of microfluidic devices has shown the promise to improve culturing scalability.^[22, 25]^ Microfluidic technologies can also provide precise spatiotemporal delivery of media and reagents, which has proved effective in controlling the local culturing environment in 2D cell cultures.^[30, 31]^ Although most microfluidic technologies for 3D organoid culture use continuous fluid perfusion for long-term media exchange, the on-chip organoids are either encapsulated in gel or located away from the flow path through which fresh media is delivered.^[20–22]^ These device configurations render the effective nutrient delivery close to the organoids to still relying on a diffusion-based mechanism. The reliance on diffusion-dominated nutrient delivery, although sufficient for 2D cell culture, may not provide sufficient nutrient supply to support healthy and uniform growth of 3D organoids growing into millimeter-scale, which is necessary for them to develop critical phenotypes for development and disease studies. In addition, perfusion operating protocols vary between different devices, making transferring and standardizing these microfluidic technologies difficult.

Here, we present an integrated platform technology to address these challenges by enabling long-term culture and live sample quality control of 3D human brain organoids. Our technology combines a continuous stirred-tank reactor (CSTR)-inspired mesofluidic bioreactor platform and a machine learning-enabled organoid quality detection method of living organoids. Guided by a novel reaction-diffusion scaling theory, our CSTR-inspired bioreactor platform provides uniform and optimal nutrient delivery for robust long-term culture while allowing for longitudinal tracking of individual organoids. Our reaction-diffusion scaling theory highlights the robustness of the CSTR-inspired bioreactor platform against different fluid perfusion protocols, which facilitates the adaptation of our technology. Using the high-quality bright-field images generated for each brain organoid during culturing, we create a classifier to select organoids with comparable structural and molecule profiles to traditional cultures, which strongly recapitulate developmental trajectories seen *in vivo*. Transcriptome analyses of organoids revealed that our mesofluidic bioreactor promoted organoid development while reducing cell death. Our method is robust and easily generalizable and thus is potentially applicable for many 3D organoid manufacturing processes.

## 2. Results

### 2.1. Design of a mesofluidic CSTR bioreactor platform for 3D organoid culture

Although there have been previous demonstrations of devices employing diffusive media exchange mechanisms, 3D organoids cultured under diffusion-dominated conditions usually cannot grow healthily into millimeter-scale due to the inefficient nutrient supply.^[20, 22–26, 32]^ To understand this culturing limitation, we compared the diffusive flux of critical nutrients (e.g., oxygen, small molecules such as GSK-3α/β inhibitors; CHIR, ALK5/TGF-β type I Inhibitors; SB, etc.) to the organoid with the nutrient consumption rate by the growing organoid. We found that in the 3D geometry, there exists a critical organoid radius over which the diffusion-dominated nutrient supply can no longer support the nutrient requirements of a growing organoid (**Figure 1a**). This critical organoid radius can be expressed as ***a_critical_*** = 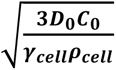, where ***D*_0_*C*_0_** characterizes the external diffusive flux of the certain nutrient and ***γ_cell_ρ_cell_*** characterizes the nutrient consumption rate of an organoid (details in Supporting Information). In particular, when oxygen is the limiting nutrient, we estimated the length scale of this critical radius to be on the order of 1 millimeter. Brain organoids typically need to be grown larger than 1 millimeter in diameter to exhibit relevant tissue structures. ^[1, 2, 26]^Therefore, microfluidic devices using only diffusive media exchange mechanisms are not adequate to be applied for 3D brain organoids.

**Figure 1.**
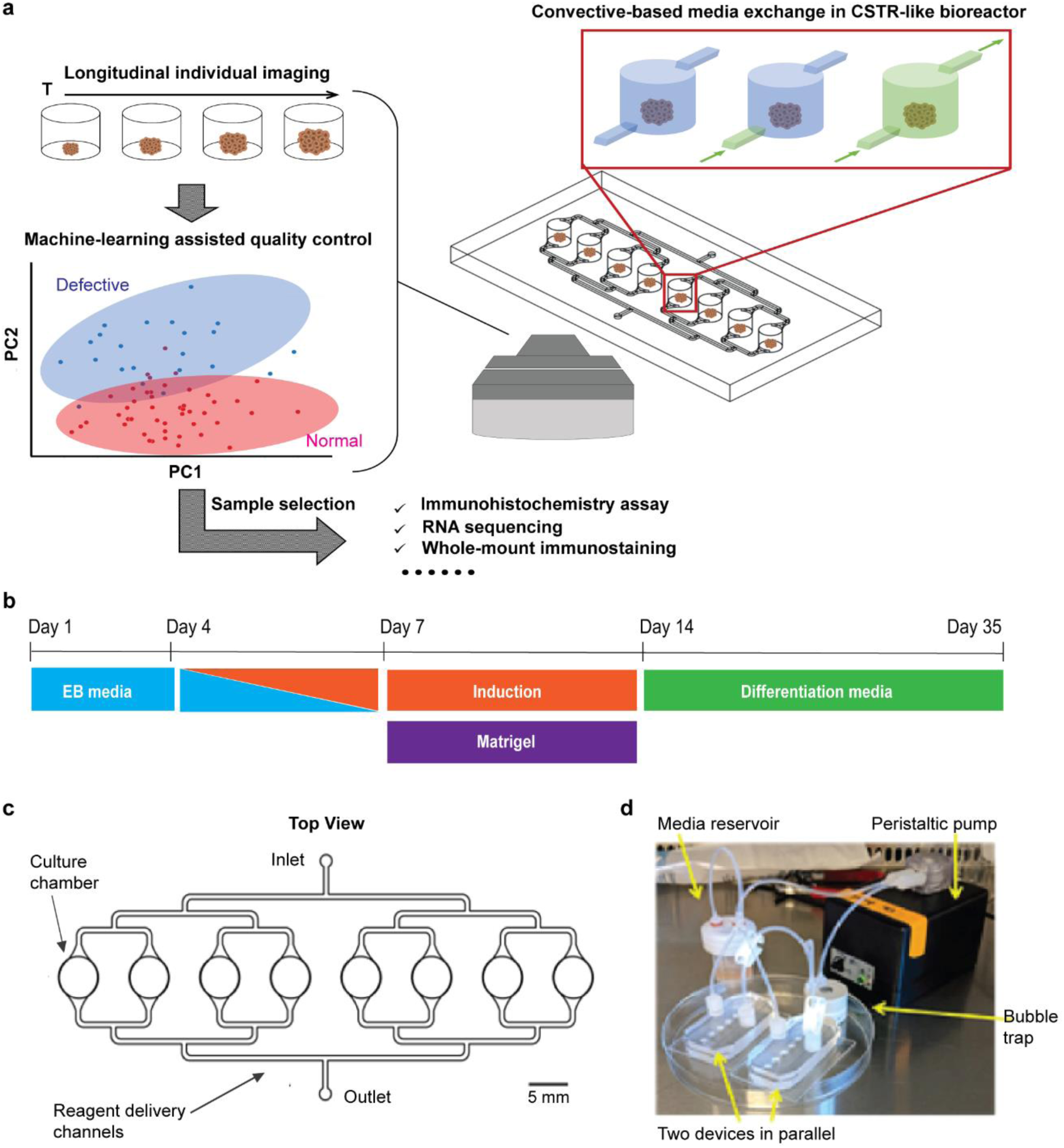
Automated mesofluidic bioreactor platform for organoid culture and analysis. (((a). Schematic of the automated pipeline for culture and non-invasive quality control of organoids. The organoids are cultured individually in an automated mesofluidic bioreactor platform. Longitudinal bright-field metrics are obtained throughout the culture process and later analyzed by a machine learning classifier model to separate organoids that develop normally from those that do not grow as expected. The organoids that underwent normal development are then analyzed using conventional molecular tools of organoid characterization. (b). Schematic diagram of forebrain organoid protocol. (c). The top view of the device design shows unique features. (d). Image describing organoid culture platform and individual parts; the peristaltic pump provides on-demand delivery of reagents to individual chambers in the culture device. Two devices are connected parallel to a media reservoir and a peristaltic pump that drives the flow. A bubble trap is located upstream of the devices to minimize bubble formation in the tubing and devices. The entire setup can be moved in and out of an incubator while maintaining sterility.))

To enable the robust long-term culture of 3D human brain organoids, we thus designed a convective-based mesofluidic bioreactor device inspired by the classic CSTR concept. Our device consists of eight culturing bioreactor chambers in parallel, which are connected to fluidic channels for perfusing media and reagents through the chambers (Figure 1a and c). The culture medium is perfused from the inlet channel at the bottom of the chamber and exits the outlet channel at the top. Although we do not agitate the media with mixers, the anti-sediment convection mechanism ensures the media is effectively well-stirred and nutrients well distributed in the chamber, which meets the CSTR criteria. The chambers are connected through a cascade of bifurcating channels to a single device inlet and outlet set, enabling consistent flow and reagent exchange between all culturing chambers (Figure 1d). A portable peristaltic pumping system controls the convective media exchange. A commercial inline bubble trap (Diba Omnifit) is placed upstream of the device to minimize the incidence of bubble generation during the multi-week-long culture.

To allow for the growth of millimeter-scale organoids, we incorporated several unique mesoscale geometrical features in our device. Each culture chamber was designed to be 5mm wide to account for the increase in organoid size as they mature. The culture chamber height was designed to be on the order of mms to minimize the fluid shear stress experienced by the cells in culture. The shear stress (T= 6uQh^-2^w^-1^) was calculated to be 0.0013 dyn cm^-2^ at the following conditions: Q=11mlhr^-1^, w=5mm, h=5mm. A bifurcated 0.6mm-wide channel connects all of the chambers; the media delivery channels are designed to expose all organoids to the same media composition. The culturing chambers were patterned to conform to the 96-well plate footprint, making our device compatible with other standard laboratory equipment such as a multichannel pipette.^[33]^

The mesoscale device was then applied for long-term culturing of 3D forebrain organoids as described by previous protocols (Figure 1b).^[1]^ We integrated our mesofluidic device with high-content imaging of individual forebrain organoids to monitor the morphology of the organoids during their development. Using the longitudinal, multi-dimensional bright-field metrics of individual organoids with machine learning tools, we achieved non-invasive quality control of organoids grown in the device to perform sample pre-selection for downstream molecular analysis (Figure 1a).

### 2.2. Sufficient nutrient delivery and robust 3D organoid culture enabled by convective-based CSTR culturing strategy

Our mesofluidic CSTR bioreactor platform promotes the healthy development of organoids by enabling uniform convective reagent exchange in individual organoid chambers (Figure 1a). High-quality bright-field images of individual organoids can be obtained longitudinally by tracking culture chambers in the mesofluidic platform (Figure 2a). We then assessed the growth of the organoids using two time-dependent metrics: the overall organoid size by its projected area (Figure 2b) and the organoid growth rate (normalized change in diameter of an organoid, Figure 2c). Under the experimental conditions, the organoids underwent a significant increase in size and growth rate during the culturing process, indicating sufficient nutrient delivery to the samples.

**Figure 2.**
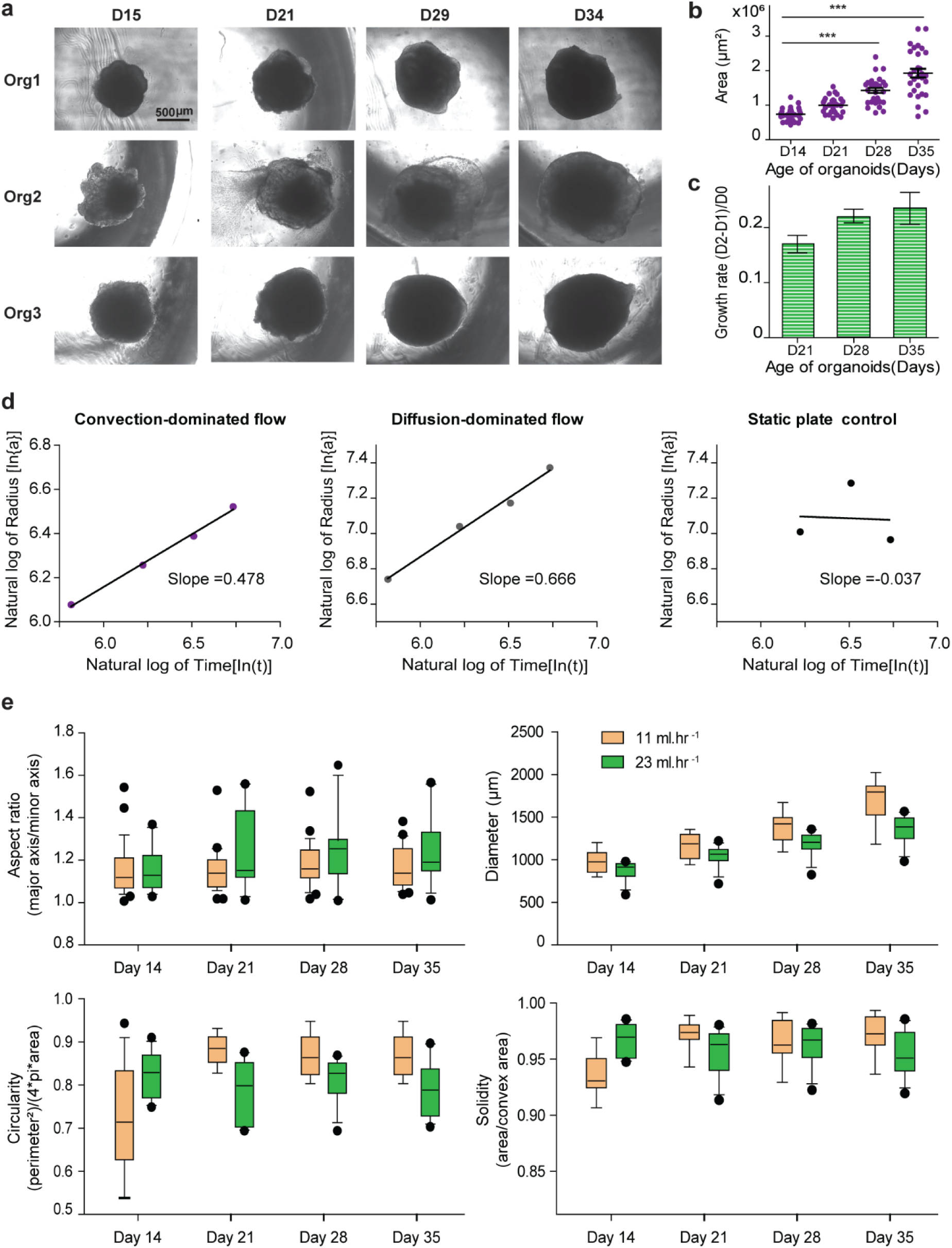
Convection-based culturing enables sufficient nutrient delivery and robust organoid culture. (((a). Representative images of organoids grown in the device. Each row represents an organoid grown in the device and each column shows changes in organoid morphology over time (b). Growth characterization of organoids grown in the device. Measuring change in area over time. Using a one-way ANOVA test with Bonferonni correction. *** indicates p-value < 0.0001. Error bars indicate S.E.M. (c). Growth characterization of organoids grown in the device. Measuring change in growth rate over time. Error bars indicate S.E.M. (d). Log-log plot of average organoid radius (indicated by a) vs. time (indicated by t) for convection-based and conventional diffusive culture methods. (e). Box-whisker plot of various bright-field metrics characterizing organoid quality in convective-based method at different flow rates (diameter, solidity, circularity, and aspect ratio). Using a two-tailed unpaired t-test with Welch correction for the two groups. N = 15 organoids for 11ml hr^-1^ condition. N = 12 organoids for 23ml hr^-1^. p-value = 0.3623, 0.5335, 0.0923, 0.9056 for diameter, circularity, aspect ratio and solidity respectively. Data is representative of four devices and two devices respectively for the 11ml hr^-1^ condition and 23 ml hr^-1^ condition. Only organoids that could be tracked for the entirety of the culture process via bright field imaging were used in this analysis. Scale bar: 500µm.))

To validate the sufficient nutrient delivery by our platform, we developed a reaction-diffusion scaling theory of organoid growth cultured in the CSTR-like mesofluidic bioreactors. Using dimensional analysis, we found that under the CSTR culturing condition, a healthy organoid growth follows a distinct square-root dynamic with respect to the culturing time. The relationship between the organoid radius *a* and the culturing time *t* can be expressed as: 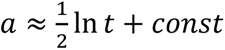 (details in Supplementary Note). Using the average radius of organoids cultured under our convective-based CSTR condition, we observed that the growth of organoids indeed exhibited this square-root scaling relationship (Figure 2d), demonstrating the consistent and sufficient nutrient supply by our platform.

In contrast, we further evaluated the growth dynamic of the organoids in two passive culturing conditions without active media exchange (Figure 2d): a diffusion-dominated condition in which the nutrient is only supplied on one side of the chamber and delivered to organoids by passive diffusion, and a static condition in which the organoids are cultured on a plate without any media agitation or mixing. In contrast to organoids cultured in our platform, the characteristic square-root scaling relationship in the organoid growth dynamic is lost for organoids cultured under these passive conditions (Figure 2d). These observations are likely due to the resulting nutrient deficiency from culturing under these conditions. Over the long culture period, the organoids are likely to undergo more significant cell death forming a larger necrotic core inside the organoid. Under this condition, the cells that continue to grow and divide are only the ones close to the edge of the organoid. These organoids can show a super 1/2 slope on the ln *a* − ln *t* plot. This super-1/2 scaling relationship is indeed observed in the middle plot in Fig. 2d (slope = 0.666). We also observed that these unhealthy organoids tend to fall apart during culture, likely a result of the larger necrotic core (data not shown). This result implies that to sustain the long-term healthy growth of organoids, the active media exchange condition offered by our convective-based CSTR mechanism is proven necessary.

It is worth noting that this scaling relationship between radius and time holds true regardless of the pumping scheme, as long as the CSTR condition is satisfied (Figure S3a). To test this outcome experimentally, we evaluated the growth and development of the organoids at different flow rates; we measured several metrics such as the organoid diameter, solidity, circularity, and aspect ratio, using the bright field images (Figure 2e).

These metrics indicate that the development of organoids in our platform is within the normal bounds. We show that at different flow rates, there are no statistical differences between these growth metrics during the culturing process (Figure 2e and Figure S4b). Our results also imply that our platform and culturing strategy are robust against fluctuations in culturing parameters such as flow rate, which is especially important in the long-term culture of brain organoids.

### 2.3. Organoids cultured in the mesofluidic bioreactor show low intra-device variability

In flow-through devices and bioreactors, transport of medium and crosstalk between cultures may influence variability among organoids. We next examined whether the location of the organoid in our platform influenced its morphological properties. Using the previously described bright-field metrics (diameter, aspect ratio, circularity, and solidity), we characterized the shape and size of organoids cultured in different on-chip chambers. When comparing the organoid diameter at the beginning (day 14) and the end (day 35) of the culture process, we observed an increase in organoid diameter over time, indicative of organoid growth in all positions. However, no statistically significant relationship was observed between the organoid position and diameter at the end of the culture process (by a Spearman correlation analysis). Our results were consistent regardless of the reagent exchange condition (**Figure S4a and Figure S5a**).

Additionally, we evaluated the effect of organoid on-chip culture position on other shape-based features, such as the organoid circularity, solidity, and aspect ratio. We observed no influence of the organoid position in the device on the shape of the organoids. Similarly, when we compared shape-based features at the beginning and end of the culture process, the same conclusions hold true: as expected, a general increase in organoid circularity and solidity and a decrease in aspect ratio can be observed between day 14 and day 35. We observed no statistically significant relationships between the organoid position and these metrics, regardless of the perfusion/reagent exchange condition (Figure S4b, c and d; Figure S5b, c and d). In total, we observed an expected organoid-to-organoid variability within a single device using the bright-field metrics in all reagent exchange conditions, but the variability is independent of the on-chip culture position of organoids in the device. Overall, our results suggest that transport in the system is sufficient to prevent depletion of nutrients supplied to the downstream organoids.

### 2.4. Non-invasive quality control of organoids enabled by high-content, longitudinal bright-field imaging

Successfully culturing an organoid that resembles the target organ in structure and function requires extensive optimization and effort.^[34]^ This is largely due to the long time required for organoid culture and the inherent heterogeneity in the self-assembly process of 3D cellular systems to form complex structures. Currently, the phenotypical heterogeneity of samples is predominantly characterized only by downstream molecular analysis steps, which limits the screening applicability of 3D organoids.^[9, 16, 35, 36]^Hence, it is critical to develop methods that quantify live morphological features, such as size and shape, to identify and select organoid cultures suitable for downstream analysis.

Here, the high-content, longitudinal imaging of individual organoids in our integrated mesofluidic platform allows us to differentiate organoids that develop normally during culture from those that do not (**Figure 3a, Figure S6a**). We found that the differences between organoids that develop normally and those that do not can be quantified using several live features obtained by bright-field images across multiple time points. Based on the images, we evaluated thirty-five features comprising of direct measures (area, circularity, solidity, etc.) and derived measures (perimeter, deviation from scaling factor, diameter, etc.) related to the organoid size and shape, as well as features related to image intensity (mean pixel intensity, standard deviation of pixel intensity) (Figure S6b, **Table S2)**. Based on these features obtained from longitudinal tracking of the organoids, we observed distinctions in these two populations of organoids grown in the mesofluidic platform (a subset showing significant differences between the two populations can be seen in Figure 3b). Some features are more predictive than others. For example, shape-based metrics (circularity, aspect ratio, etc.) can better differentiate the organoids, while metrics related to the organoid size (area, diameter, etc.) revealed little or no statistical difference between the two groups (Figure S6c). Additionally, intensity-based metrics showed statistical differences at certain time points but not consistently throughout the culture period. The inconsistency in the intensity-metrics may be attributed to variability in the imaging conditions, and do not seem to correlate with organoid health.

**Figure 3.**
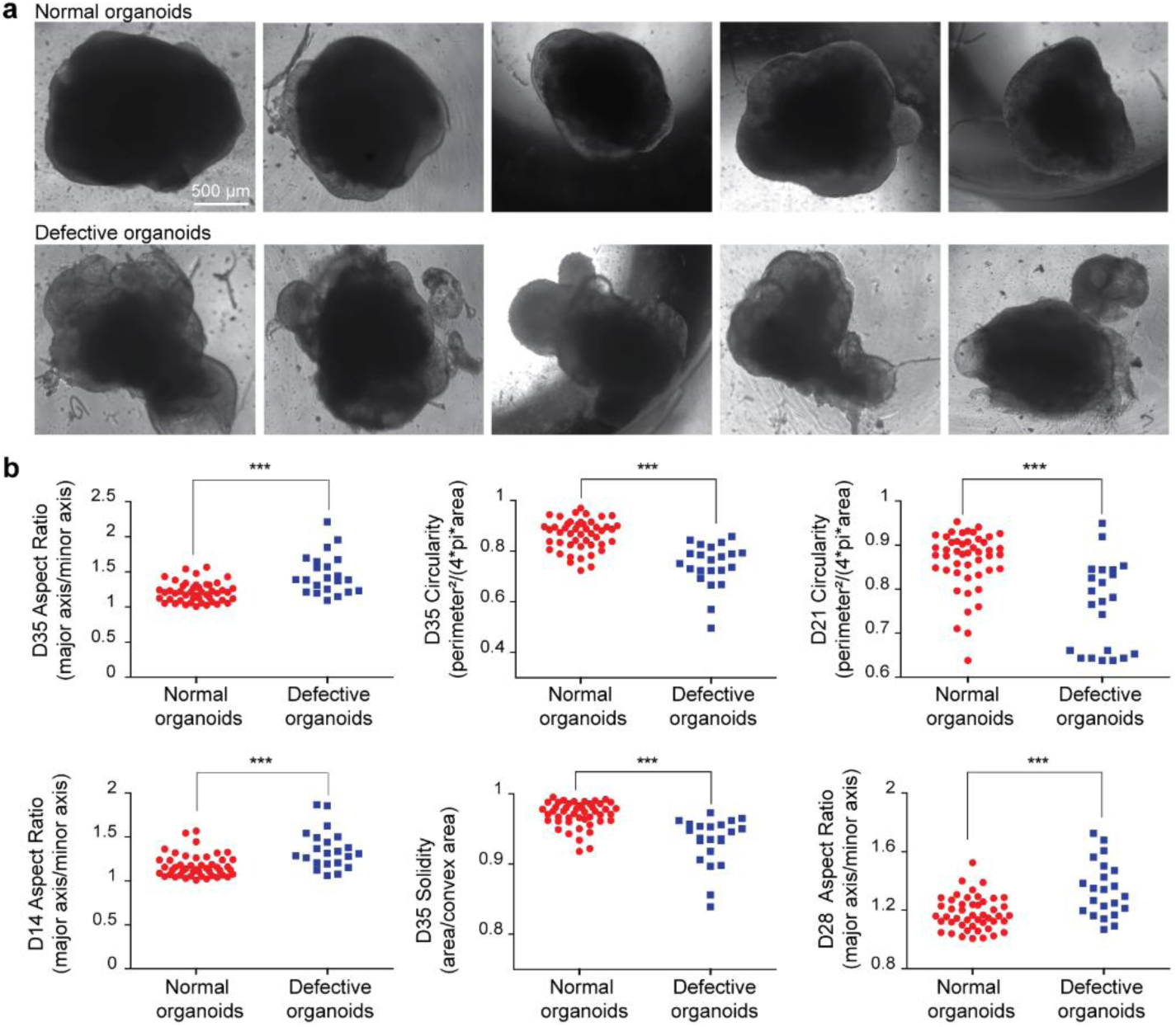
High-content brightfield images of organoids in culture can be used to determine the biological quality of organoids. (((a). Representative images of day 35 organoids that develop normally vs. defective organoids. (b). Plot showing differences in bright-field features between organoids that develop normally (red) and defective organoids (blue) in the microfluidic platform. Using a two-tailed unpaired t-test with Welch correction for the two groups. *** indicates p-value < 0.0001. Scale bar: 200μm. Data is representative of 15 devices containing 6-8 organoids obtained from all the perfusion conditions tested with the platform.))

Furthermore, when comparing individual metrics, there is a large overlap between the two groups. These results indicate that although differences exist between the two experimental groups based on bright-field metrics, one metric alone is insufficient to capture these differences. We thus employed a support vector machine (SVM) classifier to classify the physiological quality of organoids based on multiple morphological features.[37] The SVM classification successfully separates the two populations of organoids (normal organoids vs. defective organoids) (Figure 4a). Our classifier shows high specificity and accuracy in separating the groups (Figure 4b). The SVM model also identified the key features that significantly contributed to the classification. We observed that out of the thirty-five features, using a combination of twelve features was sufficient to obtain a high accuracy score (∼0.9) (Figure 4c). These twelve features were mostly shaped-related, non-derived measurements such as the organoid circularity and aspect ratio. Importantly, these twelve features spanned the entire culture period highlighting the necessity of longitudinally tracking to enable non-invasive quality control during the culture process (Figure 4d).

**Figure 4.**
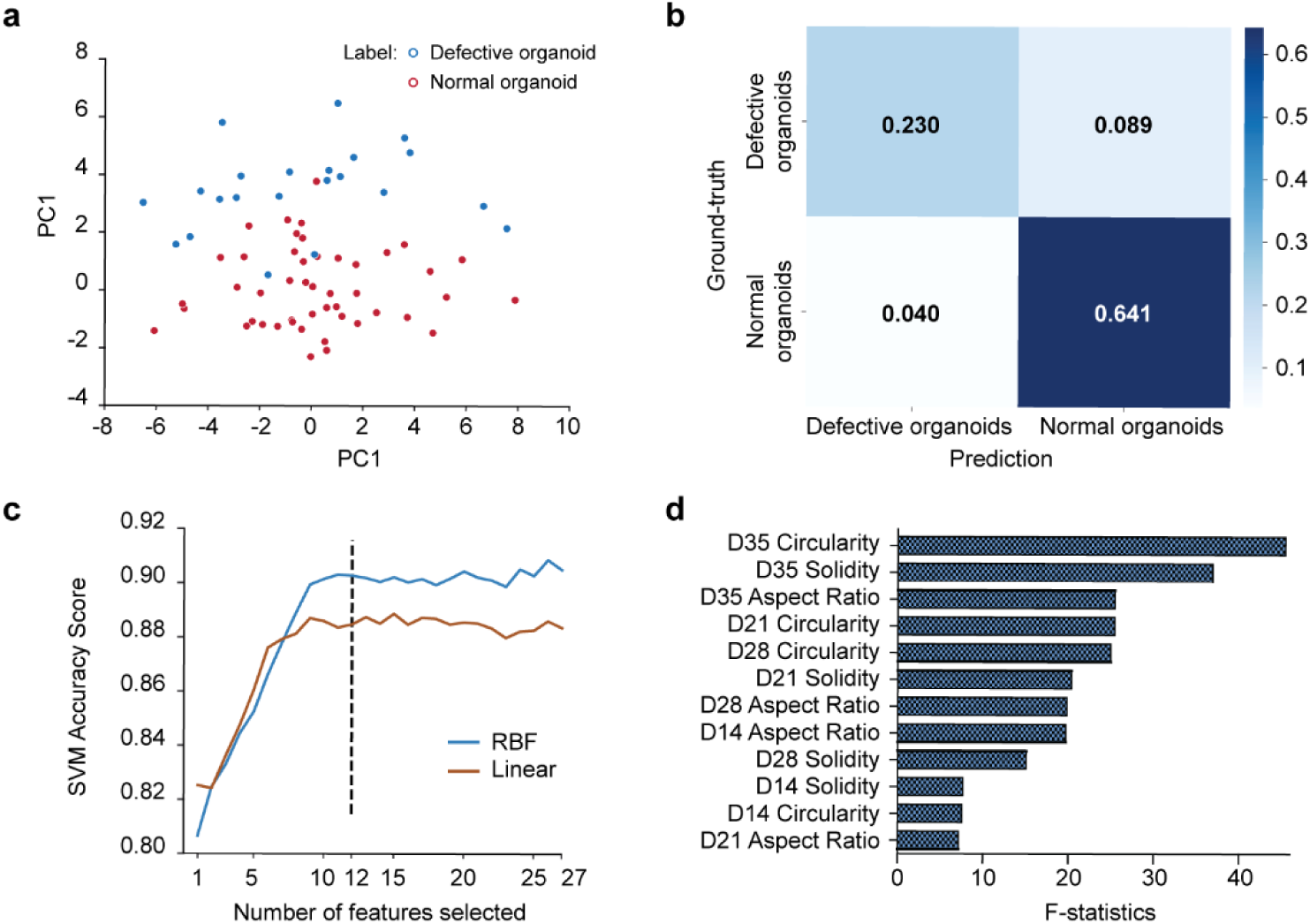
Based on the longitudinal, high-content BF images, ML-based computer vision tool can be used for the precise classification of 3D organoids based on quality. (((a). PCA plot showing the high separation between organoids based on the physiological quality & growth dynamics. (b). Heatmap showing the accuracy and specificity of the SVM classifier. (c). Plot of accuracy score vs. the number of features used in SVM classification. (d). Plot showing top 12 features used in classifier model highlighting the need for longitudinal tracking.))

We next validated the efficacy of our morphological feature-based classification by identifying differences between these two populations of organoids on the molecular scale using whole-mount immunostaining. In brain organoids, cells organize around the ventricle-like neural rosette structures as new cortical neurons migrate along radial glia fibers.^[38]^ The localization of various cell types around these rosette structures has enabled identifying and analyzing various structures and patterns present within the brain organoids.^[1, 39]^ Using whole-mount staining, we analyzed the presence and localization of several characteristic markers such as sex-determining region Y-box 2 (SOX2), microtubule-associated protein 2 (MAP2), and cortical plate markers (CTIP2) around the neural rosette structures of both populations of organoids.

Our analysis revealed that organoids identified by our classification as defective show abnormal rosette morphology, which corresponds to low expression of characteristic markers of development (Figure 5a). These organoids appeared to be less differentiated evidenced by the larger, less complex rosette structures visible on Day 35 of culture. In contrast, the normally developed organoids show more uniform rosette morphology and distribution, along with proper localization of neuronal markers (CTIP2, MAP2, TUJ1) around the rosette structures (Figure 5b and Figure S7). These organoids had smaller neural tube structures more integrated with surrounding cells, which is indicative of more mature forebrain organoid structures.^[33]^ Hence, enabled by the integrated mesofluidic platform and longitudinal tracking, our machine learning-based classification offers a real-time sorting method to select organoids based on physiological quality and growth dynamics, reducing the uncertainty in results for downstream molecular analysis. Our results also demonstrate the ability to observe organoid phenotypes in both fluorescent and non-fluorescent modes using our integrated platform.

**Figure 5.**
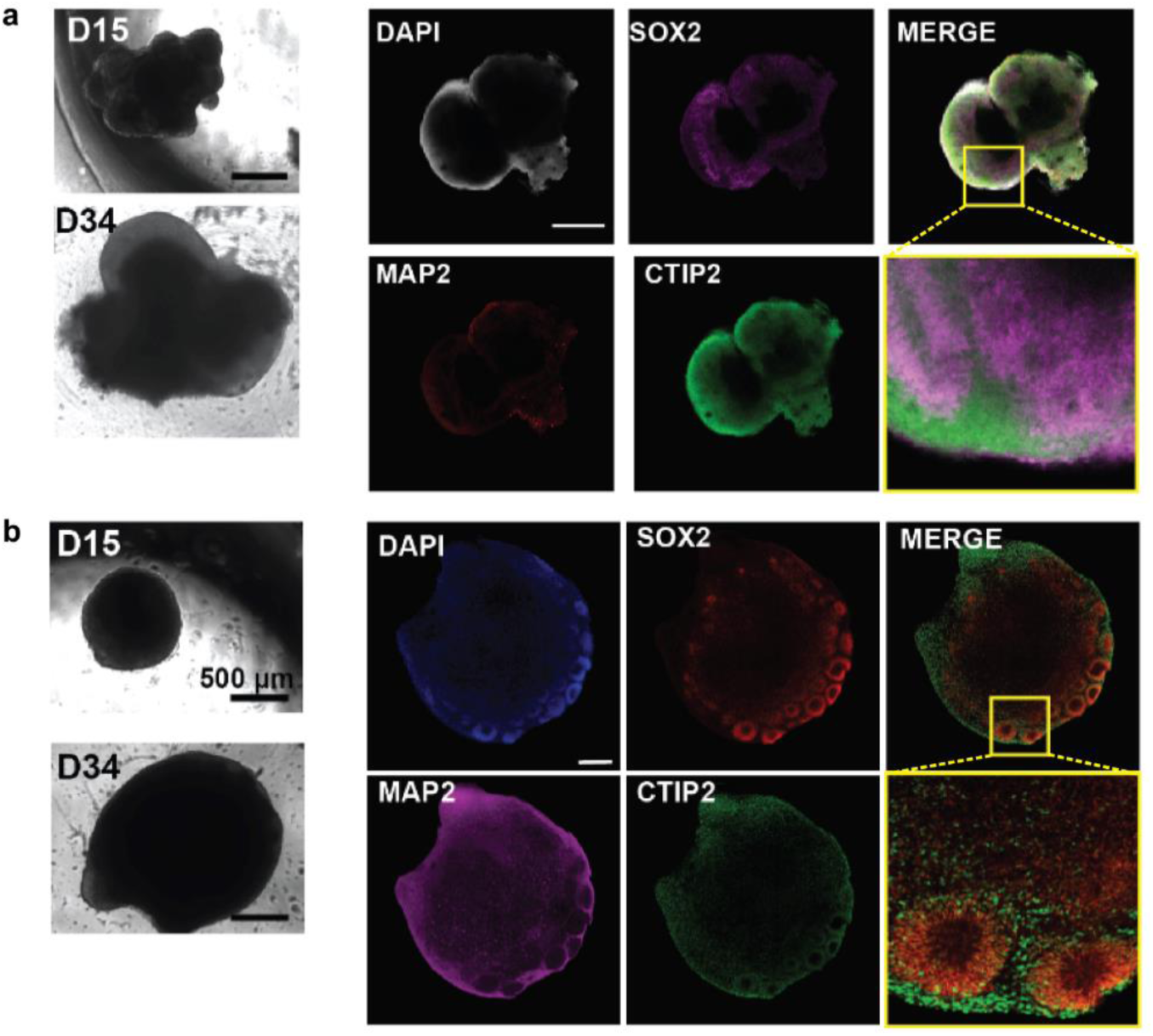
Molecular characterization of normal and defective organoids showing differences in tissue differentiation. (((a). Representative bright-field image of a defective organoid at the beginning (day14) and end (day35) of experiments accompanied by an optical slice of the same organoid showing that characteristic markers of brain development are missing or not properly localized. Insert showing zoomed-in regions around rosette structure in the organoid. Black scale bar: 500μm, White scale bar: 200μm. (b). Representative bright-field image of an organoid that normally developed at the beginning (day14) and end (day 35) of experiments accompanied by an optical slice of the same organoid showing that characteristic markers of brain development are present and properly localized. Black scale bar: 500μm, White scale bar: 200μm.))

A similar approach has been recently applied to perform sample pre-selection or sorting for drug screening applications, in which two rounds of manual and one round of automatic quality control were necessary to perform sample selection.^[13]^ Additionally, an arbitrarily defined exclusion criterion was needed for sample selection post the drug screening. Our method, in contrast, offers the advantage of incorporating longitudinal data from the culture process in the classification, potentially reducing bias in the sample pre-selection for downstream analysis.

### 2.5. Cellular characterization of 3D organoids cultured in the mesofluidic CSTR bioreactor

Using the pre-selected samples from our SVM classification, we evaluated cell types and structural features of brain development from day 28 and day 35 organoids grown in our mesofluidic CSTR bioreactor using immunohistochemistry (IHC) and compared with control organoid culture from conventional platforms (spin omega bioreactors).^[1]^ Representative images of day 28 organoids are shown in **Figures 6 a and b** (top rows: in our platform; bottom rows: in spin omega) and **Figure 6e**. Representative images of day 35 organoids are shown in Figures 6 c and d (top rows: in our platform; bottom rows: in spin omega). We first assessed the generation and proliferation of neural progenitor cells (NPCs) in organoids on days 28 and 35. As expected, organoids cultured in our platform exhibited polarized neuroepithelium-like structures resembling neural tubes containing SOX2+ NPCs (Figure 6 a-d, top rows). These neural tube structures are integrated with surrounding cells, indicative of mature forebrain organoid structures. In addition to aiding with identifying neural rosette formation, the SOX2+ areas mark the presence of ventricular zone (VZ)-like areas (Figure 6a, b and d). Neural progenitor cells migrate basally from the VZ to the SVZ.^[40]^ Hence, we next evaluated the presence of markers for cell proliferation (Ki67). Ki67 staining on day 28 indicated that the proportion of Ki67+ NPCs was lower in the device organoids compared to the control. As the organoids mature, the number of proliferating progenitor cells is expected to decrease as the SOX2+ NPCs start to exit the cell cycle. Our results indicate that the relative number of proliferating progenitor cells decreased between day 28 and day 35 in the Spin Omega (Figure 6a, c and f). The relative number of proliferating progenitor cells in the devices stayed the same between day 28 and day 35, possibly due to a decrease in the overall count of SOX2+ cells as the organoids mature.^[41]^ We also quantified the number of neural rosette structures present in the organoid samples between organoids cultured in the Spin omega and the automated platform. We observed that although the organoids in Spin omega had a higher number of neural rosettes at day 28, the number of rosettes at day 35 is comparable between organoids cultured in the two different culturing conditions (**Figure S8**). The differences in the number of neural rosettes could also contribute to the differences in the number of proliferating progenitors present in organoids cultured in both systems (Figure 6f).

**Figure 6.**
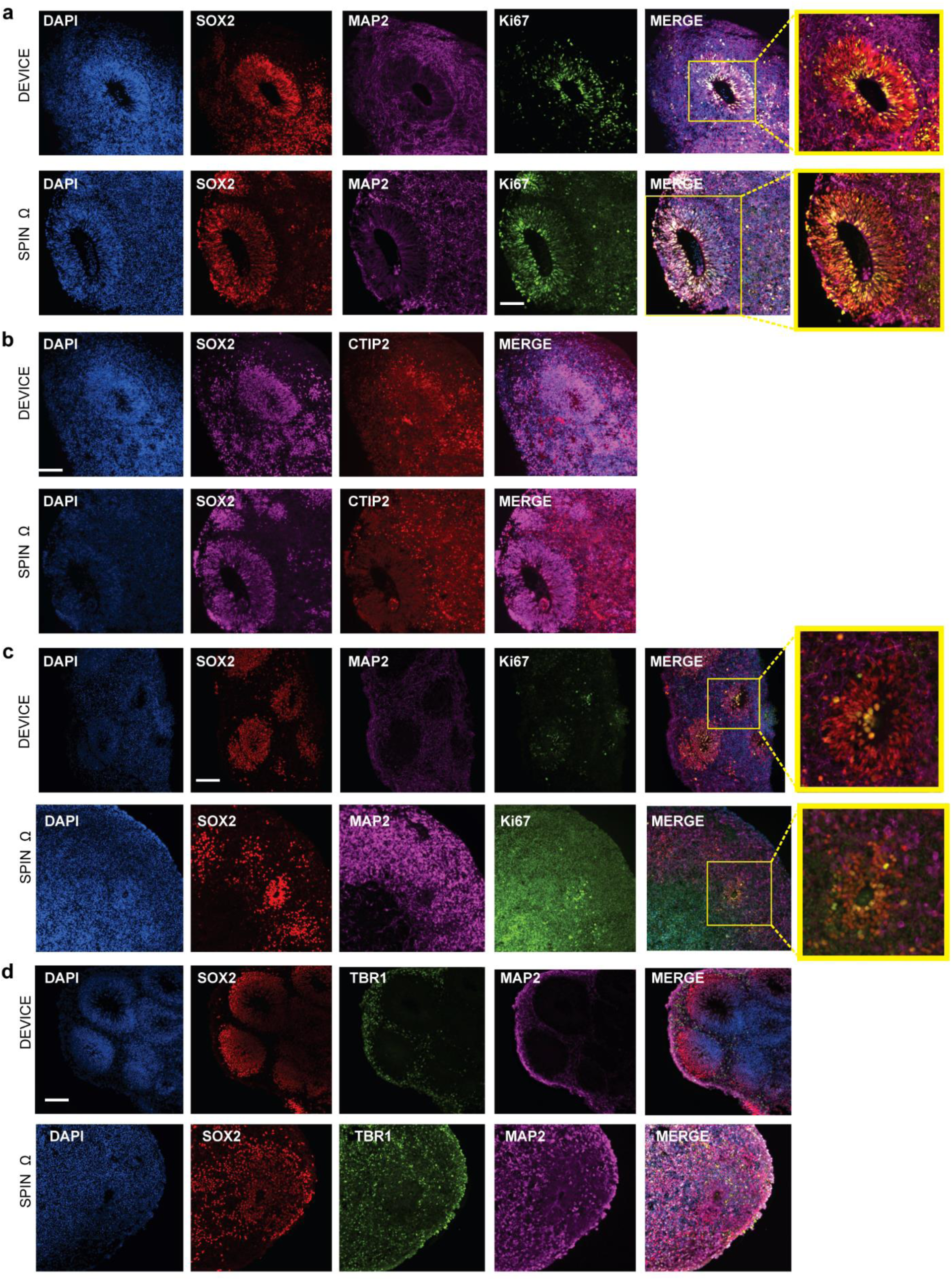
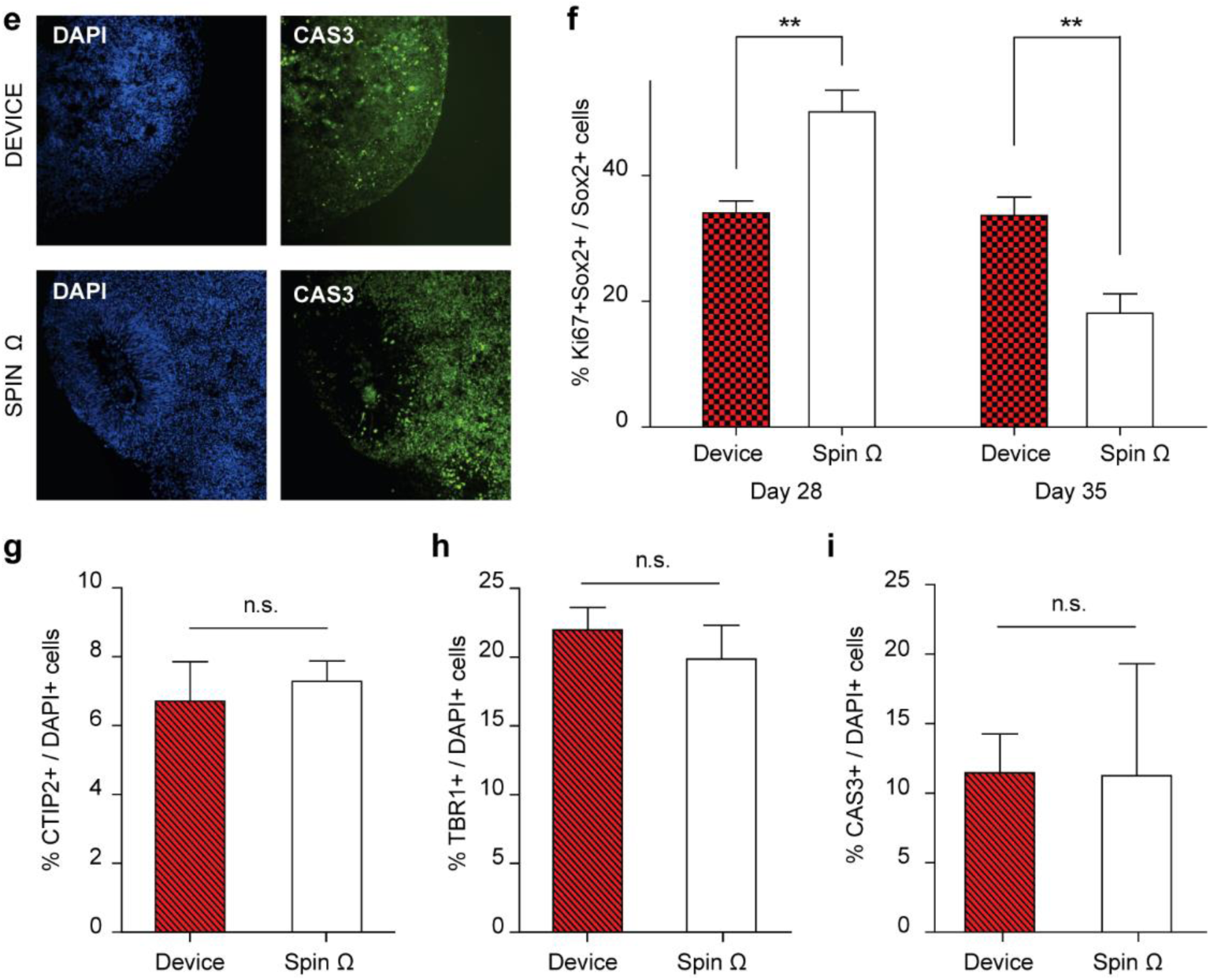
The molecular signature of the pre-selected organoids cultured in the mesofluidic platform shows normal maturation of cell phenotypes in 3D organoids. (((a). Representative immunohistochemistry (IHC) images of day 28 organoids cultured under convective-based culture method vs. spin omega showing the progression of neuronal development. Organoids were stained for a neural progenitor marker (Sox2), a proliferation marker (Ki67), and subventricular zone areas (MAP2). Scale bar: 200μm. (b). Representative immunohistochemistry (IHC) images of day 28 organoids cultured under convective-based culture method vs. spin omega showing the progression of neuronal development. Organoids were stained for a neural progenitor marker (Sox2), an intermediate progenitor marker (TBR2), and neuronal markers (CTIP2 & MAP2). Scale bar: 200μm. (c). Representative immunohistochemistry (IHC) images of day 35 organoids cultured under convective-based culture method vs. spin omega showing the progression of neuronal development. Organoids were stained for a neural progenitor marker (Sox2), a proliferation marker (Ki67), and subventricular zone areas (MAP2). Scale bar: 200μm. (d). Representative immunohistochemistry (IHC) images of day 35 organoids cultured under convective-based culture method vs. spin omega showing the progression of neuronal development. Organoids were stained for a neural progenitor marker (Sox2) and neuronal markers (TBR1 & MAP2). Scale bar: 200μm. (e). Representative immunohistochemistry (IHC) images of day 28 organoids cultured under convective-based culture method vs. spin omega showing the progression of neuronal development. Organoids were stained for a cell death marker (CASPACE 3). Scale bar: 200μm. (f-i). Quantification of results in 5a-e. Bar plots showing the percentage of proliferative progenitor cells per slice (Ki67+Sox2+/Sox2+ cells) at day 28 and day 35(f), the percentage of mature neurons out of all cells per slice (CTIP2+ cells/DAPI+ cells) at day 28 (g), the percentage of mature neurons out of all cells per slice (TBR1+cells/DAPI+ cells) at day 35 (h), and the percentage of dead cells out of all cells per slice (CASPACE3+/DAPI+ cells) at day 28(i). For all bar plots, error bars indicate S.E.M. Using a two-tailed unpaired t-test with Welch correction for the two groups. Data is representative of 6-8 organoids and 2 devices per experiment group.))

Following cell proliferation assessment, we evaluated the extent of organoid differentiation using markers of mature neurons [T-box brain 1(TBR1), CTIP2 deep layer neurons & MAP2] on day 28 and day 35. On day 28, organoids grown in our platform showed the presence of neuronal markers (MAP2) but a minimal presence of cortical plate markers (CTIP2 deep layer neurons) (Figure 6c, top row). As the organoids matured on day 35, we observed an increase in mature neurons, particularly cortical plate makers TBR1, indicating that more differentiated cells formed in the organoids over time (Figure 6d, top row). This development trend is confirmed by comparing with the control culture in spin omega bioreactors (Figure 6 b and d, bottom row; Figure 6 h and i). Finally, we assessed cell death in the 3D forebrain organoids by quantifying caspase-3 (CASP3) staining (Figure 6e). The percentage of CASP3+ cells in relation to the total number of cells (DAPI+) confirmed that all organoids cultured in our platform had low levels of cell death, comparable to the spin omega controls (Figure 6j). Overall, these results indicate that our integrated platform for organoid culturing supports the healthy growth of 3D forebrain organoids over a long period and the selection of optimal samples for downstream molecular analysis.

### 2.6. Organoids cultured in the automated platform show low inter-device variability

Given the inherent heterogeneity in the self-assembly process of 3D cellular systems to form complex structures, we evaluated if culturing in the platform contributed to increasing the variance in phenotypic outcomes of the organoids. Specifically, we quantified the variability of internal structures of the organoids across devices cultured under the same perfusion condition. We evaluated the effect of culturing in the devices on the neural rosette size, which is believed to be the site of neurogenesis in brain organoid systems. Although slight variations in the neural rosette size exist, these differences were not significantly different (**Figure S9a and b).**

We also assessed the variability in the presence of neural progenitor cells and mature neurons in the organoids. Regarding cellular identities, we assessed the variance in SOX2+, TBR2+, and CTIP2+ cell types using IHC of Day 28 organoids (Figure S9c, d, e). We observed that across devices, although there were some differences in the number of cells between devices, there was similar variability in the number of cell types in the organoid sections. Future studies will entail performing additional proteomic and transcriptomic analyses to characterize the variability across devices in our system comprehensively. Nonetheless, these preliminary results indicate that our proposed culturing method does not contribute to increasing the variability in the metrics we assessed.

### 2.7. Transcriptome analysis of organoids cultured in the automated platform

Additionally, given the observed cellular changes in organoids cultured in the mesofluidic bioreactors compared to that in the SpinΩ, we asked how the culture conditions could impact gene expression during organoid development. We performed bulk RNA-seq using organoids cultured in both mesofluidic and SpinΩ bioreactors at different developmental stages, day 28 and day 35, and identified a number of differentially expressed genes (DEGs) in the SpinΩ compared to the mesofluidic bioreactors in a developmental stage-dependent manner (**Figure 7a and d**). Gene ontology (GO) analyses of day 28 and day 35 organoids suggest that up-regulated DEGs in organoids cultured in the SpinΩ are associated with cell death, many of which have been implicated in inflammatory cell apoptotic proves, regulation of apoptotic process, regulation of neuron death, neuron apoptotic process, and activation of NF-kappaB-inducing kinase activity (Figure 7b, e, j-l); while down-regulated DEGs in organoids cultured in the SpinΩ are associated with neurodevelopment, including nervous system development, neuron development, neurogenesis, neuron differentiation, axon guidance, and neuron projection development (Figure 7c, f, g-i). To further compare forebrain organoids to *in vivo* human brain development, we performed large-scale comparisons of our organoid RNA-seq datasets with transcriptome datasets of 3 different human cortical subregions, including ventrolateral prefrontal cortex (VLPFC), orbital frontal cortex (OFC), and dorsolateral prefrontal cortex (DLPFC), at 4 developmental stages (**Figure S10**). These analyses revealed a temporal correlation between organoid and fetal human cortical development, and organoids cultured in mesofluidic bioreactors exhibited an accelerated development compared to the organoids cultured in SpinΩ (Figure S10). Overall, these results indicate that our mesofluidic platform promotes the development of organoids while reduces cell death by inhibiting the apoptotic pathway. We note that the analysis performed in this paper used organoids derived from control cell lines. To extend our method to drug screening and disease modeling applications, it is required to include organoids derived from isogenic hiPSCs lines and more control cell lines. Inclusion of additional hiPSCs lines with inserted or removed genetic mutation(s) in our method will facilitate the identification of bright-field phenotypes caused by the mutation(s) of interest from phenotypes caused by culture-related improper growth.

**Figure 7.**
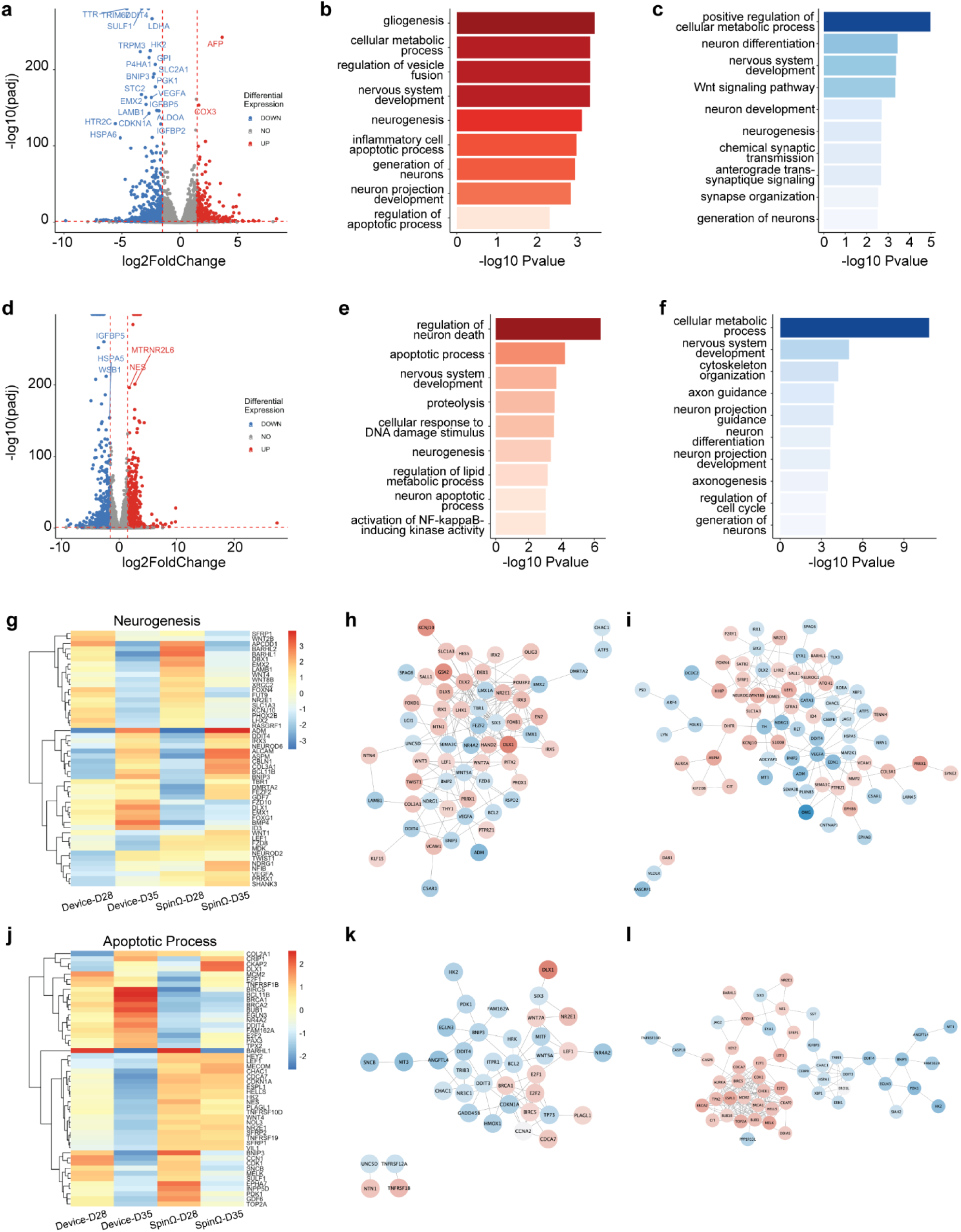
Transcriptomic comparisons of organoids cultured in the mesofluidic and SpinΩ bioreactors. (( (a). Shown is a volcano plot of gene expression change at day 28 human forebrain organoids cultured in the SpinΩ platform compared to the organoids cultured in the mesofluidic platform. Colored dots indicate statistically significant genes with false discovery rate < 0.20. Blue dots are down-regulated genes (log fold change < 0) and red dots are up-regulated genes (log fold change > 0) in SpinΩ compared to mesofluidic bioreactor. (b-c). GO Ontology overrepresentation tests using PANTHER (http://www.pantherdb.org/) for up-regulated (b) and down-regulated (c) DEGs identified from the RNA-seq of day 28 forebrain organoids cultured in SpinΩ compared to mesofluidic bioreactor is shown. P values have been adjusted for multiple testing by Bonferroni correction. (d). Shown is a volcano plot of gene expression change at day 35 human forebrain organoids cultured in the SpinΩ platform compared to the organoids cultured in the mesofluidic platform. Colored dots indicate statistically significant genes with false discovery rate < 0.20. Blue dots are down-regulated genes (log fold change < 0) and red dots are up-regulated genes (log fold change > 0) in SpinΩ compared to mesofluidic bioreactor. (e-f). Gene Ontology overrepresentation tests using PANTHER for up-regulated (e) and down-regulated (f) DEGs identified from the RNA-seq of day 35 forebrain organoids cultured in SpinΩ compared to mesofluidic bioreactor is shown. P values have been adjusted for multiple testing by Bonferroni correction. (g). Shown is a heatmap of significant DEGs enriched in neurogenesis in day 28 and day 35 forebrain organoids cultured in the SpinΩ and mesofluidic bioreactors. (h-i). Interactome plots of neurogenesis are presented. Significant DEGs found in day 28 (h) and day 35 (i) forebrain organoid RNA-seq results are shown and highlighted. Red represents up-regulated genes and blue represents down-regulated genes of organoids cultured in SpinΩ compared to mesofluidic bioreactor. (j). Shown is a heatmap of significant DEGs enriched in apoptotic process in day 28 and day 35 forebrain organoids cultured in the SpinΩ and mesofluidic bioreactors. (k-l). Interactome plots of apoptotic process are presented. Significant DEGs found in day 28 (h) and day 35 (i) forebrain organoid RNA-seq results are shown and highlighted. Red represents up-regulated genes and blue represents down-regulated genes of organoids cultured in SpinΩ compared to mesofluidic bioreactor. Data are from 2 devices per experimental group and 4-5 organoids per device.))

### 3. Conclusion

In this work, we engineered an integrated platform for long-term culture and live sample quality control of 3D human brain organoids. By analyzing the reaction-diffusion transport characteristics of a 3D multicellular system, we developed a unique mesofluidic CSTR bioreactor design to enable sufficient and robust nutrient delivery for 3D organoids. Our device thus allows for long-term culture of 3D forebrain organoids that grow beyond one millimeter and enables the development of critical tissue phenotypes. We compared the organoids cultured in our automated platform to those grown in standard spinning bioreactors. Our results indicate that the forebrain organoids develop as expected, with cell types and structures comparable to those obtained in a Spin Omega bioreactor for culture times up to 35 days. These results demonstrate that our mesofluidic CSTR bioreactor platform can replicate the effects of a miniaturized spinning bioreactor in supporting forebrain organoid differentiation.

Although previous efforts have successfully optimized differentiation protocols to reduce batch-batch differences in organoid cultures, there still exists variability at the individual organoid level that needs to be addressed or accounted for.^[17–19, 42]^ Our platform allows for the culture and monitoring of individual organoids and is compatible with various standard imaging-based methods of organoid characterization (fluorescence and non-fluorescence). Therefore, our platform enables multimodal assessment and analysis of organoids on the individual level both during and after the culture process, which could further the understanding and control of heterogeneity between individual 3D organoid systems.

We note that our goal with using SVM was to highlight the utility of longitudinal BF imaging in performing quality control of live organoids. However, we acknowledge that all image classification with conventional techniques (e.g. SVM, ADAboost, XGboost, random forest) fundamentally have to rely on arbitrarily chosen features. That is why domain-specific knowledge had been important in such processes. We can use deep learning, especially convolutional neural network (CNN) models to identify the best-working features from a set of image-label pairs without any prior knowledge, an approach that has been explored successfully prior^[34, 43]^. This approach can potentially be more efficient in performing the same tasks we demonstrated and can be explored in future projects related to this work. Furthermore, many of the parameters that are demonstrated in the paper are now very easily automatically measured. Many commercial and open-source packages that measure these parameters exist, allowing anyone without much training in ML to perform similar analyses.

Several microfluidic approaches for organoid culture exist and have made significant contributions to the field. ^[20–29]^ Nonetheless, there exists key distinctions between the proposed work and these references. Most of the cited papers do not discuss a crossflow bioreactor system to enhance organoid culture. The culturing devices in these references rely solely on passive diffusion for nutrient delivery, which may not guarantee a constant and sufficient nutrient supply near the 3D organoids, a critical aspect identified in our analysis. A more recent publication demonstrated the use of convection-based perfusion culture to promote neuronal differentiation and mitigate hypoxia and cell death in mm-scale tissue constructs.^[28]^ Although, most of the data presented is obtained from 1 week old organoids, they demonstrate the importance of perfusion during long-term culture of mm-sized tissue constructs, supporting the results presented in this paper. Our device introduces a fundamentally innovative design by facilitating a direct flow across the organoids. This unique feature is of paramount importance as it utilizes convection mechanisms to maintain adequate nutrient levels near the organoids. Furthermore, the existing work lack comprehensive characterization of flow dynamics and nutrient transport, which is essential for guiding device design. As a result, there is no assurance that the devices proposed in prior studies can be readily implemented in diverse laboratory settings, especially when addressing different research questions and necessitating distinct experimental setups. In contrast, our work provides a broadly applicable and generalizable theory for micro/meso scale bioreactor design. Additionally, the cited references do not incorporate imaging or label-free monitoring, and even when they do, the utility beyond organoid size tracking remains unclear. In our paper, we take the concept of time-lapse imaging for 3D organoids to a higher level by introducing an automated strategy for real-time monitoring, encompassing multi-dimensional phenotypes. We demonstrate the significant advantages of this approach in ensuring organoid quality before embarking on costly and time-consuming downstream molecular analyses. In summary, our non-invasive quality control pipeline was enabled by a machine-learning algorithm to assess various morphological phenotypes of the organoids. We developed a simple classification method to identify healthy 3D organoids and achieved live sample pre-selection for downstream molecular analysis. Although the organoid size has historically been used as a proxy for organoid health and growth, more recent studies have shown the limitations of using a single time-point measurement such as the organoid size in non-invasively characterizing organoid differentiation.^[42]^ Our multi-criteria image analysis method highlights the need for longitudinal monitoring and multiple morphological parameters for assessing organoid quality and potentially characterizing organoid variability and disease states.^[34, 43–48]^ Although we used 3D brain organoids as an example in this study, the live morphological features and our classification algorithm can be easily generalized to characterizing other 3D organoid models, such as intestinal organoids, colon organoids, and tumor spheroids.

## 4. Experimental Section/Methods

### Design and Fabrication of Organoid Culture Chamber

The device design was drawn in SolidWorks, and molds for the devices were made using 3D printing by the company Protolabs. The molds were printed in the material Accura SL 5530. Using the 3D printed molds, microfluidic devices were fabricated in polydimethylsiloxane (PDMS) (Dow Corning Sylgard 184, Midland, MI) by soft lithography.^[49]^ Briefly, PDMS was mixed in a 10:1 ratio of pre-polymer and crosslinker, degassed to remove air bubbles, poured on the master mold, degassed a second time to remove remaining bubbles, and cured overnight at 80°C. Following curing, PDMS devices were peeled off the master molds. The molds were not pre-treated prior to use. Additionally, creating the cross-flow in the device required two-layer PDMS fabrication. The mold for both layers was identical. For both layers of features, PDMS was poured on the mold to a height of ∼5 mm to define the height of the culture chamber. Following curing and peeling, cylindrical chambers were made in both feature layers by manually punching holes with a 5 mm biopsy punch (VWR). Inlet and outlet holes were punched with a 2 mm biopsy punch (VWR.). The bottom PDMS layer was plasma bonded to a 1mm thick glass slide. Finally, the top and bottom PDMS layers were plasma bonded together and left in an oven at 80°C overnight to strengthen the bond.

### Human iPSC culture and forebrain organoid differentiation in SpinΩ

The human iPSC line (C1-2 line) was previously generated from a skin biopsy sample of a male newborn, and has been fully characterized. ^[4, 50–55]^ Human iPSCs were cultured on irradiated MEFs in human iPSC medium consisting of D-MEM/F12 (Invitrogen), 20% Knockout Serum Replacement (KSR, Invitrogen), 1X Glutamax (Invitrogen), 1X MEM Non-essential Amino Acids (NEAA, Invitrogen), 100 µM β-Mercaptoenthanol (Invitrogen), and 10 ng ml^-1^ human basic FGF (bFGF, PeproTech) as previously described. ^[4, 47–52]^ Forebrain organoids were formed and cultured according to published protocols.^[1]^ Briefly, embryoid bodies (EBs) were first formed from hiPSC cultures by detaching the hiPSC colonies with Collagenase Type IV and culturing in ultra-low attachment 6-well plates (Corning) in media containing DMEM/F12 (Life Technologies), 20% Knockout Serum Replacement (Life Technologies), 1X Glutamax (Life

Technologies), 1X non-essential amino acids (Life Technologies), 0.1 mM 2-mercaptoethanol (Life Technologies), 2 µM A-83 (Tocris), and 2 µM Dorsomorphin (Sigma). On days 5-6, half of the media was replaced with induction media containing DMEM/F12 (Life Technologies), 1X Glutamax (Life Technologies), 1X non-essential amino acids (Life Technologies), 1X Penicillin/Streptomycin (ThermoFisher), 1X N2 Supplement (ThermoFisher), 1 µM CHIR 99021 (Cellagen Tech), and 1 µM SB-431542 (Cellagen Tech). On day 7, organoids were embedded in individual Matrigel (Corning) drops and cultured in induction media in 6-well plates until day 14. On day 14, Matrigel was mechanically dissociated from organoids by pipetting with a 5 mL serological pipette. Organoids were then transferred to microfluidic devices or low attachment tissue culture plates for the remainder of the culture. From day 14 on, organoids were cultured in differentiation media containing DMEM/F12 (Life Technologies), 1X N2 and B27 Supplements (ThermoFisher), 1X Glutamax (Life Technologies), 1X non-essential amino acids (Life Technologies), 1X Penicillin/Streptomycin (ThermoFisher), 0.1 mM 2-mercaptoethanol (Life Technologies), and 2.5 µg mL^-1^ insulin (Sigma Aldrich). For culture in spin omega, approximately 12 organoids were cultured per well with 3 mL of media. Media was exchanged every other day.

### Organoid Culture in Automated Culture Platform

Prior to each experiment device, luer fittings (Nordson Medical), the bubble trap (Cole Parmer), and tubing (1/32" I.D. silicone tubing, 1.6mm I.D. peristaltic tubing; Cole Parmer) were sterilized by autoclaving. The day before organoid loading, devices were treated with air plasma to render the PDMS hydrophilic. After treatment, devices were immediately primed with DMEM/F12 (Life Technologies) to maintain the hydrophilicity and then re-sterilized with UV light for 1 hour. Devices were then placed in a cell culture incubator overnight after replacing the DMEM/F12 with fresh DMEM/F12.

Prior to organoid loading, devices were primed with differentiation media. Organoids were loaded into individual chambers of the device by pipetting with cut 200uL tips. Following the loading of organoids into individual chambers, the devices were sealed with 3M™ Thermally Conductive Adhesive Transfer Tape (Product No:9882) to seal the culture chambers. Finally, primed tubing and fittings were connected to the device inlet and outlet. The devices were then connected to a syringe pump or peristaltic pump. The entire setup was placed in a humidified incubator (HERAcell 240i, Thermo Scientific) for culture.

### Live Imaging and Quantification

Bright-field images of devices were acquired every other day during culture using an EVOS microscope. Organoid size and shape features were quantified from images using FIJI/ImageJ. For larger organoids, multiple images of different regions of the same organoid had to be taken due to the organoids being larger than the field of view on the EVOS microscope. The images were then stitched together using a pair-wise stitching plugin on Fiji/ImageJ before quantification.^[56]^ The mean pixel intensity and standard deviation of pixel intensity were extracted from the images using a custom python script which can be found in the following Github repository: https://github.com/lu-lab/organoid-classification

### Immunohistology and Imaging

Organoids were prepared for immunohistology and imaging. Briefly, organoids were collected from microfluidic devices or spin omega and fixed in 4% Paraformaldehyde (Sigma Aldrich) in PBS for 1 hour at room temperature. Organoids were then washed in PBS and incubated in 30% sucrose overnight. Organoids were embedded in an optimal cutting temperature compound (OCT) and sectioned with a cryostat. For immunostaining, slides were permeabilized with 0.2% Triton-X for 1 hour and then washed with blocking buffer containing 10% donkey serum and 0.1% Tween-20 in PBS for 30 minutes. Next, slices were incubated in primary antibodies in blocking buffer overnight at 4°C. Slices were washed with PBS and incubated with secondary antibodies in blocking buffer for 1 hour. Slices were then washed in PBS and incubated with DAPI. Images were collected on an epifluorescent microscope (Nikon Eclipse Ti-E microscope). Quantification of imaging data was performed using a customized Python code.

### Immunohistology Quantification

We used a custom computer vision script for the cell segmentation. The script uses the image for the nuclear stain (DAPI, Hoechst, etc.) as the input and the range of sizes of cells to be detected (For our analysis, we used a minimum cell radius of 5 and a maximum cell radius of 25). The script then creates a general outline of the organoid slice by performing a closing operation with kernel size equal to 2 maximum cell diameters. Next, it takes the original image and performs iterative adaptive thresholding with window sizes ranging from maximum cell diameter to the minimum image dimension. Each time, it adds newly identified foreground pixels to the total mask image ("bitwise or" operation). The user determines the number of steps in this process. Next, the original nuclei-stained image is passed to the Laplacian of the Gaussian blob detection filter. Local maxima of the filtered image are then detected, each designating the center of a cell. All cells detected outside the general outline of the organoid are considered artifacts and are filtered out. The remaining cell centers are next used as seeds in a watershed algorithm with boundaries set to the total mask of the organoid slice. In the next steps, the intensity of each cell is calculated in each of the given channel images using the cell outlines identified during the segmentation.

We used several approaches to determine positively and negatively stained cells in each channel: k-means clustering with k=2, otsu, triangle, and mean thresholding. The choice of the right thresholding method is dependent on numerous factors (imaging conditions, quality of the antibodies used for staining, etc.) and is left to the discretion of the user. For the SOX2, Ki67, and TBR1 channels, we performed k-means clustering to separate the positively and negatively stained cells. We performed triangle thresholding for the CTIP2 and TBR2 channels to separate the positively and negatively stained cells.

### Support Vector Machine (SVM) Classifier Model

We analyzed a total of 70 images of organoids to test our SVM model. We used the scikit-learn toolbox to implement the SVM organoid classification model and extract feature importance. ^[57]^ We employed the grid-search over a hyperparameter grid for both RBF kernel SVM and linear SVM with 5-fold cross-validation. The grid search was performed 1000 times, and a hyperparameter set that peaked in repetitions most frequently was chosen for the further analyses for both RBF kernel SVM (gamma=0.001, C=10) and linear SVM (C=1). A summary of the results from the grid-search is shown in **Table S3**. In each repetition, 70% of the dataset was randomly selected as the training set and used in the grid-search process, and the left 30% was used for testing the optimal model from the grid-search. A confusion matrix was obtained by averaging the testing result over the repetitions**. Figure S11** provides a summary of SVM’s performance on both the training and test data set as we gradually increased the size of the training data set. This was used to determine the number of images required to minimize overfitting of the SVM model.

### Feature selection by F-statistics

For the feature selection, we carried out an ANOVA analysis. We varied the number of features to use from 1 to 27 and measured the accuracy of the classification model with a given feature number. For a given number k as the number of features, we first selected top k features having high F-statistics in the ANOVA analysis.

Then we split the dataset into the 70% training set and the 30% test set. The RBF kernel SVM and linear SVM with the best hyperparameter sets obtained above are fitted to the training set and tested to the test set to get the accuracy score. The steps are repeated 1000 times to get an average score for the given k. The link to the Github repository for the code can be found here: https://github.com/lu-lab/organoid-classification

### Whole-Mount Immunostaining of Organoids Using iDISCO

A modified iDISCO protocol was used for staining and clearing the whole organoids.^[58]^ Glycine was not added to the iDISCO permeabilization solution, and the PTwH solution was substituted with PBST. The organoid samples were fixed with 4% PFA. for 60 minutes at 4 °C. Following fixation, the samples were washed three times with 1X PBS. The samples underwent a methanol / H_2_O dehydration series (33%,66%) for 1 hour each at room temperature. Next, the samples were incubated overnight in 66% DCM/33% methanol at room temperature. Next, the samples were bleached in chilled fresh 5% H_2_O_2_ in methanol overnight at 4°C. The samples were then washed in 100% methanol and then chilled at 4 °C for 1 hour. The samples were then subjected to a rehydration series with methanol / H_2_O (66%, 33%) for 1 hour each at room temperature. This was followed by a 2x wash in PTx.2 (PBS 10x, 0.2% wt Triton X-100) for 1 hour each. The samples were incubated in Permeabilization solution (PTx.2, DMSO) at 37 °C overnight, followed by blocking (PTx.2, Donkey Serum, DMSO, NaCl, 5% BSA) for 1 hour at 37 °C prior to incubation with the primary antibody staining solution at 4°C for 3 days. This was followed by four washing steps in PBST (10X PBS, DI water, Tween 20) for 1 hour each at room temperature. The samples were incubated in secondary antibody staining solution at 4°C for 3 days, followed by 3 washing steps in PBST. The samples were then stained with TOPRO-3 (Thermofisher, 1:1000) for 1 hour at room temperature, followed by another three washes in PBST for 20 minutes each at room temperature. The samples were subjected to a methanol / H2O dehydration series (33%,66%, 100%) for 1 hour each at room temperature, followed by a 2-hour incubation in 66% DCM / 33% methanol at room temperature. The samples were then incubated in 100% DCM to wash the methanol. This was followed by incubation in dibenzyl ether (DBE) prior to imaging in the mesofluidic platform with a Zeiss 700A confocal microscope using an L.D. 20x objective (N.A. = 0.4).

### Whole-Mount Immunostaining of Organoids Using BABB

Organoids were recovered from the mesofluidic device prior to staining. Staining protocol was conducted using pre-established protocols.^[59]^ Briefly, the organoid samples were fixed with 4% PFA for 60 minutes at 4°C. Following fixation, the samples were washed three times with 1X PBS. This was followed by a 2-hour incubation in Dent’s Bleach [MeOH:DMSO:H2O2 (4:1:1)] to promote antibody penetration and quench auto-fluorescence. The samples were washed in 100% methanol for 10-15 minutes at room temperature. The samples were then equilibrated to PBS through a MeOH dehydration series (75%, 50%, 25%) for 10 minutes each at room temperature prior to blocking in 0.5% TNB [0.1 M Tris-HCl, pH 7.5, 0.15 M NaCl, 0.5% (w/v) blocking reagent (PerkinElmer FP1020)] for 2 hours at Room temperature. Post blocking, the samples were incubated in primary antibodies diluted in 0.5% TNB overnight on a rotating rack at 4°C. The following day, the samples were washed five times for 1 hour each with PBS 1x at room temperature. The samples were then incubated in secondary antibodies diluted in 0.5% TNB overnight at 4°C. The following day, the samples were washed three times in PBS 1x for 20 minutes each at room temperature before applying the nuclear stain.

The samples were washed again with PBS 3 times for 20 minutes each prior to undergoing a methanol dehydration series (25%, 50%, 75% in PBS 1x) for 20 minutes each. After complete equilibration with 100% methanol, the organoid samples were transferred to an Attofluor imaging chamber and a BABB [benzyl alcohol: benzyl benzoate(1:2)]: methanol solution (1:1) was added to the wells for 5 minutes. After clearing the samples, the BABB solution was removed and replaced with fresh BABB before imaging them using a laser scanning confocal microscope (Zeiss 700 A, L.D. 20x objective).

### RNA Isolation, RNA-seq Library Preparation, and Sequencing

Human forebrain organoids were homogenized in Trizol (Invitrogen) and processed according to the manufacturer’s instructions. RNA-seq libraries were generated from 1 μg of total RNA from duplicated or triplicated samples per condition using the TruSeq LT RNA Library Preparation Kit v2 (Illumina) following the manufacturer’s protocol. An Agilent 2100 BioAnalyzer and DNA1000 kit (Agilent) were used to quantify amplified cDNA and to control the quality of the libraries. A qPCR-based KAPA library quantification kit (KAPA Biosystems) was applied to accurately quantify library concentration. Illumina HiSeq2500 were used to perform 150-cycle PE sequencing. Raw reads were examined for quality issues to ensure library generation and sequencing were suitable for further analysis. RNA-seq reads were trimmed using Trimmomatic Version 0.40 and aligned using Salmon v1.9.0.^[60, 61]^ Significantly differentially expressed genes were identified using DESeq2 by comparing regularized log transformed counts between samples with adjusted p-values < 0.05.^[62]^ Gene Ontology analyses for Biological Processes were performed using GOstats.^[63]^ Human dorsolateral prefrontal cortex RNA-seq datasets from different life stages were obtained from BioProject PRJNA245228.^[64]^ RNA-seq gene expression for different time points of fetal development and different brain regions were obtained from the Allen Brain Atlas (www.brain-map.org). R programming language was used to perform all analyses and to produce all figures except protein interaction networks which were generated using Cytoscape.^[65]^

### Statistical information

Experiments with replicate data are represented as the mean +/- standard error. The corresponding figure legends detailed sample sizes (N) and the number of independent experiments. Statistical analyses were performed using GraphPad Prism software. Statistical tests were performed using either a two-tailed Mann-Whitney U test (with Welch’s correction) or non-parametric one-way ANOVA (with Kruskal-Wallis) or two-way ANOVA combined with Dunn’s multiple comparison’s test or Bonferonni multiple comparison’s for comparison of individual samples. P values less than 0.05 were considered significant.

## Acknowledgements

We acknowledge the National Institutes of Health, National Science Foundation, Simons Foundation, and Marcus Foundation (NIH R01NS096581, R01GM088333, and R21EB021676, NSF 0939511, 1764406, and 1648035, Simons Foundation, Marcus Center for Therapeutic Cell Characterization and Manufacturing grants to H.L.) for funding, A. Aiyar, S. Szabo, C. Martin, M. Luong, A. Shaw for technical assistance, C. Sedlock and D. Franta (3M™ company) for the generous donation of the adhesive tapes used for sealing the microfluidic devices, and C. Luthio for moral support. S. Charles, E. Jackson-Holmes and G. Sun contributed equally to this work.

## References

1. Qian, X.; Nguyen, Ha N.; Song, Mingxi M.; Hadiono, C.; Ogden, Sarah C.; Hammack, C.; Yao, B.; Hamersky, Gregory R.; Jacob, F.; Zhong, C.; Yoon, K.-j.; Jeang, W.; Lin, L.; Li, Y.; Thakor, J.; Berg, Daniel A.; Zhang, C.; Kang, E.; Chickering, M.; Nauen, D.; Ho, C.-Y.; Wen, Z.; Christian, Kimberly M.; Shi, P.-Y.; Maher, Brady J.; Wu, H.; Jin, P.; Tang, H.; Song, H.; Ming, G.-l., Cell 2016, 165 (5), 1238–1254. DOI 10.1016/j.cell.2016.04.032.

2. Lancaster, M. A.; Renner, M.; Martin, C.-A.; Wenzel, D.; Bicknell, L. S.; Hurles, M. E.; Homfray, T.; Penninger, J. M.; Jackson, A. P.; Knoblich, J. A., Nature 2013, 501 (7467), 373–379. DOI 10.1038/nature12517.

3. Mariani, J.; Coppola, G.; Zhang, P.; Abyzov, A.; Provini, L.; Tomasini, L.; Amenduni, M.; Szekely, A.; Palejev, D.; Wilson, M.; Gerstein, M.; Grigorenko, E. L.; Chawarska, K.; Pelphrey, K. A.; Howe, J. R.; Vaccarino, F. M., Cell 2015, 162 (2), 375–390. DOI 10.1016/J.CELL.2015.06.034.

4. Kang, Y.; Zhou, Y.; Li, Y.; Han, Y.; Xu, J.; Niu, W.; Li, Z.; Liu, S.; Feng, H.; Huang, W.; Duan, R.; Xu, T.; Raj, N.; Zhang, F.; Dou, J.; Xu, C.; Wu, H.; Bassell, G. J.; Warren, S. T.; Allen, E. G.; Jin, P.; Wen, Z., Nature Neuroscience 2021, 24 (10), 1377–1391. DOI 10.1038/s41593-021-00913-6.

5. Jo, J.; Xiao, Y.; Sun, A. X.; Cukuroglu, E.; Tran, H. D.; Göke, J.; Tan, Z. Y.; Saw, T. Y.; Tan, C. P.; Lokman, H.; Lee, Y.; Kim, D.; Ko, H. S.; Kim, S. O.; Park, J. H.; Cho, N. J.; Hyde, T. M.; Kleinman, J. E.; Shin, J. H.; Weinberger, D. R.; Tan, E. K.; Je, H. S.; Ng, H. H., Cell Stem Cell 2016, 19 (2), 248–257. DOI 10.1016/J.STEM.2016.07.005.

6. Paşca, A. M.; Sloan, S. A.; Clarke, L. E.; Tian, Y.; Makinson, C. D.; Huber, N.; Kim, C. H.; Park, J.-Y.; O’Rourke, N. A.; Nguyen, K. D.; Smith, S. J.; Huguenard, J. R.; Geschwind, D. H.; Barres, B. A.; Paşca, S. P., Nature Methods 2015, 12 (7), 671–678. DOI 10.1038/nmeth.3415.

7. Sloan, S. A.; Darmanis, S.; Huber, N.; Khan, T. A.; Birey, F.; Caneda, C.; Reimer, R.; Quake, S. R.; Barres, B. A.; Paşca, S. P., Neuron 2017, 95 (4), 779–790.e6. DOI 10.1016/J.NEURON.2017.07.035.

8. Gabriel, E.; Wason, A.; Ramani, A.; Gooi, L. M.; Keller, P.; Pozniakovsky, A.; Poser, I.; Noack, F.; Telugu, N. S.; Calegari, F.; Šarić, T.; Hescheler, J.; Hyman, A. A.; Gottardo, M.; Callaini, G.; Alkuraya, F. S.; Gopalakrishnan, J., The EMBO Journal 2016, 35 (8), 803–819. DOI 10.15252/embj.201593679.

9. Raja, W. K.; Mungenast, A. E.; Lin, Y.-T.; Ko, T.; Abdurrob, F.; Seo, J.; Tsai, L.-H., PLOS ONE 2016, 11 (9), e0161969.

10. Seo, J.; Kritskiy, O.; Watson, L. A.; Barker, S. J.; Dey, D.; Raja, W. K.; Lin, Y.-T.; Ko, T.; Cho, S.; Penney, J.; Silva, M. C.; Sheridan, S. D.; Lucente, D.; Gusella, J. F.; Dickerson, B. C.; Haggarty, S. J.; Tsai, L.-H., The Journal of Neuroscience 2017, 37 (41), 9917–9917. DOI 10.1523/JNEUROSCI.0621-17.2017.

11. Wang, P.; Mokhtari, R.; Pedrosa, E.; Kirschenbaum, M.; Bayrak, C.; Zheng, D.; Lachman, H. M., Molecular Autism 2017, 8 (1), 11–11. DOI 10.1186/s13229-017-0124-1.

12. Groveman, B. R.; Ferreira, N. C.; Foliaki, S. T.; Walters, R. O.; Winkler, C. W.; Race, B.; Hughson, A. G.; Zanusso, G.; Haigh, C. L., Scientific Reports 2021, 11 (1), 5165–5165. DOI 10.1038/s41598-021-84689-6.

13. Park, J.-C.; Jang, S.-Y.; Lee, D.; Lee, J.; Kang, U.; Chang, H.; Kim, H. J.; Han, S.-H.; Seo, J.; Choi, M.; Lee, D. Y.; Byun, M. S.; Yi, D.; Cho, K.-H.; Mook-Jung, I., Nature communications 2021, 12 (1), 280–280. DOI 10.1038/s41467-020-20440-5.

14. A. Durens, M.; Nestor, J.; Williams, M.; Herold, K.; Niescier, R. F.; Lunden, J. W.; Phillips, W.; Lin, Y. C.; Dykxhoorn, D. M.; Nestor, M. W., Journal of Neuroscience Methods 2020, 335, 108627–108627. DOI 10.1016/J.JNEUMETH.2020.108627.

15. Suong, D. N. A.; Imamura, K.; Inoue, I.; Kabai, R.; Sakamoto, S.; Okumura, T.; Kato, Y.; Kondo, T.; Yada, Y.; Klein, W. L.; Watanabe, A.; Inoue, H., Communications Biology 2021, 4 (1), 1213–1213. DOI 10.1038/s42003-021-02719-5.

16. Quadrato, G.; Nguyen, T.; Macosko, E. Z.; Sherwood, J. L.; Min Yang, S.; Berger, D. R.; Maria, N.; Scholvin, J.; Goldman, M.; Kinney, J. P.; Boyden, E. S.; Lichtman, J. W.; Williams, Z. M.; McCarroll, S. A.; Arlotta, P., Nature 2017, 545 (7652), 48–53. DOI 10.1038/nature22047.

17. Velasco, S.; Kedaigle, A. J.; Simmons, S. K.; Nash, A.; Rocha, M.; Quadrato, G.; Paulsen, B.; Nguyen, L.; Adiconis, X.; Regev, A.; Levin, J. Z.; Arlotta, P., Nature 2019, 570 (7762), 523–527. DOI 10.1038/s41586-019-1289-x.

18. Cederquist, G. Y.; Asciolla, J. J.; Tchieu, J.; Walsh, R. M.; Cornacchia, D.; Resh, M. D.; Studer, L., Nature Biotechnology 2019, 37 (4), 436–444. DOI 10.1038/s41587-019-0085-3.

19. Fiorenzano, A.; Sozzi, E.; Birtele, M.; Kajtez, J.; Giacomoni, J.; Nilsson, F.; Bruzelius, A.; Sharma, Y.; Zhang, Y.; Mattsson, B.; Emnéus, J.; Ottosson, D. R.; Storm, P.; Parmar, M., Nature Communications 2021, 12 (1), 7302–7302. DOI 10.1038/s41467-021-27464-5.

20. Wang, Y.; Wang, L.; Zhu, Y.; Qin, J., Lab on a chip 2018, 18 (6), 851–860. DOI 10.1039/c7lc01084b.

21. Berger, E.; Magliaro, C.; Paczia, N.; Monzel, A. S.; Antony, P.; Linster, C. L.; Bolognin, S.; Ahluwalia, A.; Schwamborn, J. C., Lab on a Chip 2018, 18 (20), 3172–3183. DOI 10.1039/C8LC00206A.

22. Zhu, Y.; Wang, L.; Yu, H.; Yin, F.; Wang, Y.; Liu, H.; Jiang, L.; Qin, J., Lab on a Chip 2017, 17 (17), 2941–2950. DOI 10.1039/C7LC00682A.

23. Cai, H.; Ao, Z.; Wu, Z.; Song, S.; Mackie, K.; Guo, F., Lab on a Chip 2021, 21 (11), 2194–2205. DOI 10.1039/D1LC00145K.

24. Khan, I.; Prabhakar, A.; Delepine, C.; Tsang, H.; Pham, V.; Sur, M., Biomicrofluidics 2021, 15, 024105–024105. DOI 10.1063/5.0041027.

25. Ao, Z.; Cai, H.; Havert, D. J.; Wu, Z.; Gong, Z.; Beggs, J. M.; Mackie, K.; Guo, F., Analytical Chemistry 2020, 92 (6), 4630–4638. DOI 10.1021/acs.analchem.0c00205.

26. Cho, A.-N.; Jin, Y.; An, Y.; Kim, J.; Choi, Y. S.; Lee, J. S.; Kim, J.; Choi, W.-Y.; Koo, D.-J.; Yu, W.; Chang, G.-E.; Kim, D.-Y.; Jo, S.-H.; Kim, J.; Kim, S.-Y.; Kim, Y.-G.; Kim, J. Y.; Choi, N.; Cheong, E.; Kim, Y.-J.; Je, H. S.; Kang, H.-C.; Cho, S.-W., Nature Communications 2021, 12 (1), 4730–4730. DOI 10.1038/s41467-021-24775-5.

27. Seiler, S. T.; Mantalas, G. L.; Selberg, J.; Cordero, S.; Torres-Montoya, S.; Baudin, P. V.; Ly, V. T.; Amend, F.; Tran, L.; Hoffman, R. N.; Rolandi, M.; Green, R. E.; Haussler, D.; Salama, S. R.; Teodorescu, M., Scientific Reports 2022, 12 (1), 20173. DOI 10.1038/s41598-022-20096-9.

28. Grebenyuk, S.; Abdel Fattah, A. R.; Kumar, M.; Toprakhisar, B.; Rustandi, G.; Vananroye, A.; Salmon, I.; Verfaillie, C.; Grillo, M.; Ranga, A., Nature Communications 2023, 14 (1), 193. DOI 10.1038/s41467-022-35619-1.

29. Zhu, Y.; Zhang, X.; Sun, L.; Wang, Y.; Zhao, Y., Advanced Materials 2023, 35 (14), 2210083. DOI 10.1002/adma.202210083.

30. Demers, C. J.; Soundararajan, P.; Chennampally, P.; Cox, G. A.; Briscoe, J.; Collins, S. D.; Smith, R. L., Development 2016, 143 (11), 1884–1892. DOI 10.1242/dev.126847.

31. Manfrin, A.; Tabata, Y.; Paquet, E. R.; Vuaridel, A. R.; Rivest, F. R.; Naef, F.; Lutolf, M. P., Nature Methods 2019, 16 (7), 640–648. DOI 10.1038/s41592-019-0455-2.

32. Wang, Y.; Wang, L.; Guo, Y.; Zhu, Y.; Qin, J., RSC Advances 2018, 8, 1677–1685. DOI 10.1039/C7RA11714K.

33. Jackson-Holmes, E. L., Microfluidics-based tools for culture and multi-functional assessments of three-dimensional pluripotent stem cell derived tissues. Altanta, 2018.

34. Kassis, T.; Hernandez-Gordillo, V.; Langer, R.; Griffith, L. G., Scientific Reports 2019, 9 (1), 12479–12479. DOI 10.1038/s41598-019-48874-y.

35. Sakaguchi, H.; Kadoshima, T.; Soen, M.; Narii, N.; Ishida, Y.; Ohgushi, M.; Takahashi, J.; Eiraku, M.; Sasai, Y., Nature Communications 2015, 6 (1), 8896–8896. DOI 10.1038/ncomms9896.

36. Monzel, A. S.; Smits, L. M.; Hemmer, K.; Hachi, S.; Moreno, E. L.; van Wuellen, T.; Jarazo, J.; Walter, J.; Brüggemann, I.; Boussaad, I.; Berger, E.; Fleming, R. M. T.; Bolognin, S.; Schwamborn, J. C., Stem Cell Reports 2017, 8 (5), 1144–1154. DOI 10.1016/j.stemcr.2017.03.010.

37. Boser, B. E.; Guyon, I. M.; Vapnik, V. N. In A Training Algorithm for Optimal Margin Classifiers, Proceedings of the Fifth Annual Workshop on Computational Learning Theory, New York, NY, USA, Association for Computing Machinery: New York, NY, USA, 1992; pp 144–152.

38. Albanese, A.; Swaney, J. M.; Yun, D. H.; Evans, N. B.; Antonucci, J. M.; Velasco, S.; Sohn, C. H.; Arlotta, P.; Gehrke, L.; Chung, K., Scientific Reports 2020, 10 (1), 21487–21487. DOI 10.1038/s41598-020-78130-7.

39. Watanabe, M.; Buth, J. E.; Vishlaghi, N.; de la Torre-Ubieta, L.; Taxidis, J.; Khakh, B. S.; Coppola, G.; Pearson, C. A.; Yamauchi, K.; Gong, D.; Dai, X.; Damoiseaux, R.; Aliyari, R.; Liebscher, S.; Schenke-Layland, K.; Caneda, C.; Huang, E. J.; Zhang, Y.; Cheng, G.; Geschwind, D. H.; Golshani, P.; Sun, R.; Novitch, B. G., Cell Reports 2017, 21 (2), 517–532. DOI 10.1016/J.CELREP.2017.09.047.

40. Johnson, C. A.; Ghashghaei, H. T., Development (Cambridge, England) 2020, 147 (4), dev186056–dev186056. DOI 10.1242/dev.186056.

41. Graham, V.; Khudyakov, J.; Ellis, P.; Pevny, L., Neuron 2003, 39 (5), 749–765. DOI 10.1016/S0896-6273(03)00497-5.

42. Gehling, K.; Parekh, S.; Schneider, F.; Kirchner, M.; Kondylis, V.; Nikopoulou, C.; Tessarz, P., bioRxiv 2021, 2021.11.22.469588–2021.11.22.469588. DOI 10.1101/2021.11.22.469588.

43. Borten, M. A.; Bajikar, S. S.; Sasaki, N.; Clevers, H.; Janes, K. A., Scientific Reports 2018, 8 (1), 5319–5319. DOI 10.1038/s41598-017-18815-8.

44. Broutier, L.; Mastrogiovanni, G.; Verstegen, M. M. A.; Francies, H. E.; Gavarró, L. M.; Bradshaw, C. R.; Allen, G. E.; Arnes-Benito, R.; Sidorova, O.; Gaspersz, M. P.; Georgakopoulos, N.; Koo, B.-K.; Dietmann, S.; Davies, S. E.; Praseedom, R. K.; Lieshout, R.; Ijzermans, J. N. M.; Wigmore, S. J.; Saeb-Parsy, K.; Garnett, M. J.; van der Laan, L. J. W.; Huch, M., Nature Medicine 2017, 23 (12), 1424–1435. DOI 10.1038/nm.4438.

45. B. N, O. S.; Fleur, W.; K, D. K.; M, M. C.; Sovann, K.; Erik, v. W.; Luuk, S.; Louisa, H.; J, V. D.; Joris, v. d. H.; Warner, P.; Petur, S.; Daphne, v. d. V.; Michelle, K.; Myriam, C.; Henk, ; Monique, v. L.; J, B. H.; V, B. L.; Lodewyk, W.; Edwin, C.; Hans, C.; E, V. E., Science Translational Medicine 2019, 11 (513), eaay2574–eaay2574. DOI 10.1126/scitranslmed.aay2574.

46. Beato, F.; Reverón, D.; Dezsi, K. B.; Ortiz, A.; Johnson, J. O.; Chen, D.-T.; Ali, K.; Yoder, S. J.; Jeong, D.; Malafa, M.; Hodul, P.; Jiang, K.; Centeno, B. A.; Abdalah, M. A.; Balasi, J. A.; Tassielli, A. F.; Sarcar, B.; Teer, J. K.; DeNicola, G. M.; Permuth, J. B.; Fleming, J. B., Laboratory Investigation 2021, 101 (2), 204–217. DOI 10.1038/s41374-020-00494-1.

47. Tiriac, H.; Belleau, P.; Engle, D. D.; Plenker, D.; Deschênes, A.; Somerville, T. D. D.; Froeling, F. E. M.; Burkhart, R. A.; Denroche, R. E.; Jang, G.-H.; Miyabayashi, K.; Young, C. M.; Patel, H.; Ma, M.; LaComb, J. F.; Palmaira, R. L. D.; Javed, A. A.; Huynh, J. C.; Johnson, M.; Arora, K.; Robine, N.; Shah, M.; Sanghvi, R.; Goetz, A. B.; Lowder, C. Y.; Martello, L.; Driehuis, E.; LeComte, N.; Askan, G.; Iacobuzio-Donahue, C. A.; Clevers, H.; Wood, L. D.; Hruban, R. H.; Thompson, E.; Aguirre, A. J.; Wolpin, B. M.; Sasson, A.; Kim, J.; Wu, M.; Bucobo, J. C.; Allen, P.; Sejpal, D. V.; Nealon, W.; Sullivan, J. D.; Winter, J. M.; Gimotty, P. A.; Grem, J. L.; DiMaio, D. J.; Buscaglia, J. M.; Grandgenett, P. M.; Brody, J. R.; Hollingsworth, M. A.; O’Kane, G. M.; Notta, F.; Kim, E.; Crawford, J. M.; Devoe, C.; Ocean, A.; Wolfgang, C. L.; Yu, K. H.; Li, E.; Vakoc, C. R.; Hubert, B.; Fischer, S. E.; Wilson, J. M.; Moffitt, R.; Knox, J.; Krasnitz, A.; Gallinger, S.; Tuveson, D. A., Cancer Discovery 2018, 8 (9), 1112–1129. DOI 10.1158/2159-8290.CD-18-0349.

48. Sachs, N.; de Ligt, J.; Kopper, O.; Gogola, E.; Bounova, G.; Weeber, F.; Balgobind, A. V.; Wind, K.; Gracanin, A.; Begthel, H.; Korving, J.; van Boxtel, R.; Duarte, A. A.; Lelieveld, D.; van Hoeck, A.; Ernst, R. F.; Blokzijl, F.; Nijman, I. J.; Hoogstraat, M.; van de Ven, M.; Egan, D. A.; Zinzalla, V.; Moll, J.; Boj, S. F.; Voest, E. E.; Wessels, L.; van Diest, P. J.; Rottenberg, S.; Vries, R. G. J.; Cuppen, E.; Clevers, H., Cell 2018, 172 (1), 373–386.e10. DOI 10.1016/j.cell.2017.11.010.

49. Duffy, D. C.; McDonald, J. C.; Schueller, O. J.; Whitesides, G. M., Anal. Chem. 1998, 70, 4974–4974.

50. Wen, Z.; Nguyen, H. N.; Guo, Z.; Lalli, M. A.; Wang, X.; Su, Y.; Kim, N.-S.; Yoon, K.-J.; Shin, J.; Zhang, C.; Makri, G.; Nauen, D.; Yu, H.; Guzman, E.; Chiang, C.-H.; Yoritomo, N.; Kaibuchi, K.; Zou, J.; Christian, K. M.; Cheng, L.; Ross, C. A.; Margolis, R. L.; Chen, G.; Kosik, K. S.; Song, H.; Ming, G.-l., Nature 2014, 515 (7527), 414–418. DOI 10.1038/nature13716.

51. Kim, N.-S.; Wen, Z.; Liu, J.; Zhou, Y.; Guo, Z.; Xu, C.; Lin, Y.-T.; Yoon, K.-J.; Park, J.; Cho, M.; Kim, M.; Wang, X.; Yu, H.; Sakamuru, S.; Christian, K. M.; Hsu, K.-s.; Xia, M.; Li, W.; Ross, C. A.; Margolis, R. L.; Lu, X.-Y.; Song, H.; Ming, G.-l., Nature Communications 2021, 12 (1), 1398. DOI 10.1038/s41467-021-21713-3.

52. Bentea, E.; Depasquale, E. A. K.; O’Donovan, S. M.; Sullivan, C. R.; Simmons, M.; Meador-Woodruff, J. H.; Zhou, Y.; Xu, C.; Bai, B.; Peng, J.; Song, H.; Ming, G.-l.; Meller, J.; Wen, Z.; McCullumsmith, R. E., Molecular Omics 2019, 15 (3), 173–188. DOI 10.1039/C8MO00173A.

53. C. Xu, M.; Lee, E. M.; Wen, Z.; Cheng, Y.; Huang, W.-K.; Qian, X.; Tcw, J.; Kouznetsova, J.; Ogden, S. C.; Hammack, C.; Jacob, F.; Nguyen, H. N.; Itkin, M.; Hanna, C.; Shinn, P.; Allen, ; Michael, S. G.; Simeonov, A.; Huang, W.; Christian, K. M.; Goate, A.; Brennand, K. J.; Huang, R.; Xia, M.; Ming, G.-l.; Zheng, W.; Song, H.; Tang, H., Nature Medicine 2016, 22 (10), 1101–1107. DOI 10.1038/nm.4184.

54. Tang, H.; Hammack, C.; Ogden, Sarah C.; Wen, Z.; Qian, X.; Li, Y.; Yao, B.; Shin, J.; Zhang, F.; Lee, Emily M.; Christian, Kimberly M.; Didier, Ruth A.; Jin, P.; Song, H.; Ming, G.-l., Cell Stem Cell 2016, 18 (5), 587–590. DOI 10.1016/j.stem.2016.02.016.

55. Inglis, G. A. S.; Zhou, Y.; Patterson, D. G.; Scharer, C. D.; Han, Y.; Boss, J. M.; Wen, Z.; Escayg, A., Human Molecular Genetics 2020, 29 (15), 2579–2595. DOI 10.1093/hmg/ddaa150.

56. Preibisch, S.; Saalfeld, S.; Tomancak, P., Bioinformatics 2009, 25 (11), 1463–1465. DOI 10.1093/bioinformatics/btp184.

57. Pedregosa, F.; Varoquaux, G.; Gramfort, A.; Michel, V.; Thirion, B.; Grisel, O.; Blondel, M.; Prettenhofer, P.; Weiss, R.; Dubourg, V.; Vanderplas, J.; Passos, A.; Cournapeau, D.; Brucher, M.; Perrot, M.; Duchesnay, É., J. Mach. Learn. Res. 2011, 12 (null), 2825–2830.

58. Renier, N.; Wu, Z.; Simon, David J.; Yang, J.; Ariel, P.; Tessier-Lavigne, M., Cell 2014, 159 (4), 896–910. DOI 10.1016/j.cell.2014.10.010.

59. Ahnfelt-Rønne, J.; Jørgensen, M. C.; Hald, J.; Madsen, O. D.; Serup, P.; Hecksher-Sørensen, J., Journal of Histochemistry & Cytochemistry 2007, 55 (9), 925–930. DOI 10.1369/jhc.7A7226.2007.

60. Bolger, A. M.; Lohse, M.; Usadel, B., Bioinformatics 2014, 30 (15), 2114–2120. DOI 10.1093/bioinformatics/btu170.

61. Patro, R.; Duggal, G.; Love, M. I.; Irizarry, R. A.; Kingsford, C., Nature Methods 2017, 14 (4), 417–419. DOI 10.1038/nmeth.4197.

62. Love, M. I.; Huber, W.; Anders, S., Genome Biology 2014, 15 (12), 550. DOI 10.1186/s13059-014-0550-8.

63. Falcon, S.; Gentleman, R., Bioinformatics 2007, 23 (2), 257–258. DOI 10.1093/bioinformatics/btl567.

64. Jaffe, A. E.; Shin, J.; Collado-Torres, L.; Leek, J. T.; Tao, R.; Li, C.; Gao, Y.; Jia, Y.; Maher, B. J.; Hyde, T. M.; Kleinman, J. E.; Weinberger, D. R., Nature neuroscience 2015, 18 (1), 154–161. DOI 10.1038/nn.3898.

65. Shannon, P.; Markiel, A.; Ozier, O.; Baliga, N. S.; Wang, J. T.; Ramage, D.; Amin, N.; Schwikowski, B.; Ideker, T., Genome Research 2003, 13 (11), 2498–2504. DOI 10.1101/gr.1239303.

