## Supplemental Information for "Non-Invasive Quality Control of Organoid Cultures Using Mesofluidic CSTR Bioreactors and High-Content Imaging"

### 1. Supplementary Note. Diffusion-Reaction Model for Organoid Growth Dynamic under Microfluidic Culturing

In this paper, we use diffusion-reaction transport models and dimensional analysis to understand the growth dynamic of spherical organoids cultured in different conditions. We demonstrate two findings that provide insights to the design of microfluidic culturing condition and the interpretation of experimental organoid culturing data. First, under pure diffusive culturing condition, a nutrient depletion boundary layer forms around the growing organoid, which prevent the nutrient delivery to the organoid. As a result, there exists a critical organoid radius beyond which the diffusive condition cannot sustain the healthy growth of the organoid. Second, using microfluidic perfusion culturing method, the depletion layer can be removed to maintain a maximum nutrient supply. Under microfluidic perfusion culturing, a square root scaling is found between the organoid radius and culturing time. This consistent scaling relationship is determined by the intra-organoid diffusion-reaction dynamic and can be used to analyze the experimental data.

### 2. Lists of Terms and Symbols:

$\boldsymbol{a}$: the radius of the organoid; Unit: **m**.

$\boldsymbol{\rho_{cell}}$: the cell density in the organoid; Unit: **# of cell m$^{-3}$**.

$\boldsymbol{\gamma_{cell}}$: the per cell consumption rate of a critical nutrient; Unit: **mol s$^{-1}$ (per cell)**. *we use mole as the basic unit for nutrient.

$\boldsymbol{\eta_{cell}}$: the total amount of nutrient required to grow a new cell (for example, by cell division); Unit: **mol (per cell)**.

$\boldsymbol{C_0}$: the concentration of a critical nutrient in the fresh culturing media; Unit: **mol m$^{-3}$**.

$\boldsymbol{D_0}$: the diffusivity of the critical nutrient in the media; Unit: **m$^2$ s$^{-1}$**.

$D_{organoid}$: the diffusivity of the critical nutrient in the organoid; Unit: $\mathbf{m^2\ s^{-1}}$.

**3. The model set-up**

*We assume an organoid as a homogeneous, isotropic, spherical "reactor".* The radius of an organoid is denoted by $\boldsymbol{a}$. The organoid is composed by cells. For simplicity, we assume all cells are the same. The cell density in the organoid is denoted by $\boldsymbol{\rho_{cell}}$. A cell, once made, consumes nutrient at a consistent rate. We denote the per cell consumption rate of a critical nutrient by $\boldsymbol{\gamma_{cell}}$. *Note that the term "cell" in our model analysis is not equivalent to a real biological cell. We simply treat a "cell" as a unit volume that consumes nutrient at a constant rate.*

We consider the problem in the following analysis in the spherical coordinate. The center of the organoid is located at the origin. We are interested in how a critical nutrient (glucose, oxygen, etc.) is transported to and in the organoid and are consumed by the organoid. Therefore, we denote the concentration of such nutrient in the fresh media by $\boldsymbol{C_0}$.

**4. Analysis of organoid growth under diffusive culturing by an extra-organoid diffusion-reaction model**

In this section, we are interested in why diffusive culturing conditions fails to grow organoid over a certain size in experiments. Hence, we sought to ask a simple question: **can the nutrient supply towards the organoid by diffusive flux alone be enough to compensate the consumption by the organoid?**

To answer this question, we compare two quantities: the overall nutrient consumption rate by the organoid, and the nutrient diffusive flux towards the organoid. The overall nutrient consumption rate, $\boldsymbol{\Gamma_{organoid}}$, can be expressed as:

$$\boldsymbol{\Gamma_{organoid} = \gamma_{cell}\rho_{cell}V_{organoid} = \gamma_{cell}\rho_{cell}\frac{4}{3}\pi a^3} \qquad \text{(S1)}$$

$\boldsymbol{\Gamma_{organoid}}$ has a unit of $\mathbf{mol\ s^{-1}}$. Note that the overall consumption rate is scaled with the cube of radius of the organoid, $\boldsymbol{a^3}$.

To obtain the expression for the nutrient diffusive flux into the organoid, we consider the extra-organoid diffusive transport of such nutrient, which can be described by Laplace equation:

$$\boldsymbol{D_0\nabla^2 C^{out} = D_0\frac{1}{r^2}\frac{d}{dr}\left(r^2\frac{dC^{out}}{dr}\right) = 0} \qquad \text{(S2)}$$

where $\boldsymbol{C^{out}}$ denotes the extra-organoid concentration profile. The angular components of the spherical Laplacian operator are omitted by the isotropic assumption.

The boundary conditions can be specified by Dirichlet boundary conditions outside the organoid:

$$\begin{cases} \boldsymbol{C^{out}(r = \infty) = C_0} \\ \boldsymbol{C^{out}(r = a) = C_1} \end{cases} \quad \text{(S3)}$$

where $\boldsymbol{r = \infty}$ describe the far-field condition in which the media is fresh; $\boldsymbol{r = a}$ describes the concentration in the vicinity of the organoid.

In order to promote the diffusive nutrient flux, a depleted boundary layer in the vicinity of the organoid must be established. The maximum inward flux is achieved when the nutrient is completely depleted near the organoid. Therefore, to compare the organoid consumption rate with the maximum nutrient supply possible, we render $\boldsymbol{C_1 = 0}$.

Solving the Laplace equation, we obtain the nutrient distribution outside the organoid:

$$\boldsymbol{C^{out}(r) = C_0\left(1 - \frac{a}{r}\right)} \quad \text{(S4)}$$

Therefore, the maximum integrated diffusive flux to supply nutrient towards the organoid, $\boldsymbol{J_{diffusion}}$, can be obtained as:

$$\boldsymbol{J_{diffusion} = \oiint_S D_0 \nabla C^{out}(a)\, dS = 4\pi D_0 C_0 a} \quad \text{(S5)}$$

$\boldsymbol{S}$ denotes the surface area of the organoid; $\boldsymbol{J_{diffusion}}$ has a unit of **mol s$^{-1}$**. Note that the flux is scaled with the radius of the organoid, $\boldsymbol{a}$.

Comparing the overall nutrient consumption rate $\boldsymbol{\Gamma_{organoid}}$ and the maximum diffusive nutrient flux $\boldsymbol{J_{diffusion}}$, we notice that a critical radius exists. Beyond this critical radius, $\boldsymbol{J_{diffusion}}$ is always smaller than $\boldsymbol{\Gamma_{organoid}}$, which means the nutrient supply is no longer sufficient to sustain the consumption of the organoid. We can obtain this critical radius by solving $\boldsymbol{\Gamma_{organoid} = J_{diffusion}}$, with respect to $\boldsymbol{a}$. Therefore, the critical radius is:

$$\boldsymbol{a_{critical} = \sqrt{\frac{3D_0C_0}{\gamma_{cell}\rho_{cell}}}} \quad \text{(S6)}$$

This critical radius represents the maximum viable size of spherical organoid growth and has also been described or estimated differently in previous studies of 3D spherical growth under diffusive conditions.[1-4] However, a major assumption of this analysis is that the nutrient consumption rate of the organoid is constant. Given that the organoid is composed of a wide diversity of cell types with varying levels of metabolic activity, we estimated the range of critical size values that would be supported under diffusive conditions. For our analysis, we focused on estimating the critical size values for two essential nutrients (glucose and oxygen) using parameters obtained from literature. A summary of the parameters used in this study is provided below in **Table S1.**

**Table S1. Summary of parameters used in sensitivity analysis for glucose and oxygen**

| Species | $D_0$ [$m^2$ $s^{-1}$] | $C_0$ [mM] | $\gamma_{cell}$ [mol $cell^{-1}$ $s^{-1}$] | Ref |
|---|---|---|---|---|
| Oxygen | $1\text{-}3*10^{-9}$ | 0.2 | $2.3*10^{-18}\text{-}7*10^{-16}$ | [1, 5] |
| Glucose | $6\text{-}7*10^{-10}$ | 4-55 | $9*10^{-17}\text{-}2.2*10^{-16}$ | [1, 6, 7] |

We estimated the critical organoid size at varying glucose consumption rates. Figure S1 shows the results of our analysis. As expected, the estimated critical size is lower for cells with higher nutrient consumption rates and vice versa. Additionally, the critical size is on the order of hundreds of microns to millimeters. The analysis was performed using an initial concentration ($C_0$) value of 20.4mM; a glucose concentration previously used in brain organoid cultures. [8] The glucose consumption ranges chosen were from $9*10^{-17}$mol $cell^{-1}$ $s^{-1}$ (estimated average human neuron consumption) to $2.2*10^{-16}$ mol $cell^{-1}$ $s^{-1}$ (estimated average human cortical neuron consumption).[6, 7]

**Figure S1. Sensitivity analysis of critical organoid size under diffusive conditions at different glucose consumption rates**

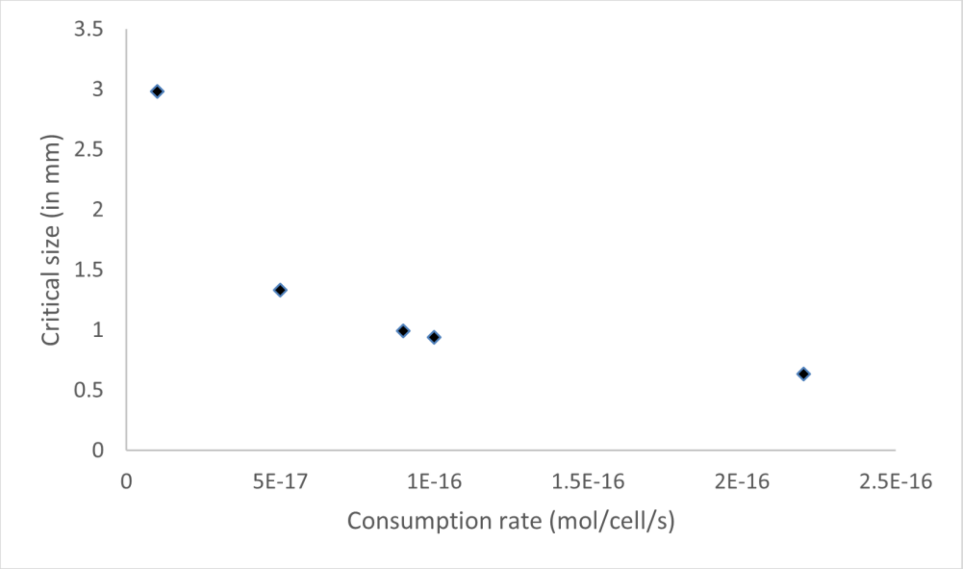

A similar analysis was performed for oxygen (Figure S2). The analysis was performed using an initial concentration ($C_0$) value of 0.2mM. The oxygen consumption ranges chosen where from 2.3*$10^{-18}$ mol $cell^{-1}s^{-1}$ (estimated hiPSC oxygen consumption) to 7*$10^{-16}$ mol $cell^{-1}$ $s^{-1}$ (estimated average human neuron oxygen consumption). [6, 7] In both oxygen and glucose analysis, the estimated organoid density ($\rho_{cell}$) used was on the order of $10^{14}$ cell $m^{-3}$ as determined by previous studies.[1, 8]

We observed a similar trend between the critical size and the nutrient consumption rate when performing the analysis for oxygen. However, oxygen appears to be more limiting than glucose due to the lower critical sizes calculated. These results are consistent with previous analytical models of tissue construct growth under diffusive conditions.

**Figure S2. Sensitivity analysis of critical organoid size under diffusive conditions at different oxygen consumption rates**

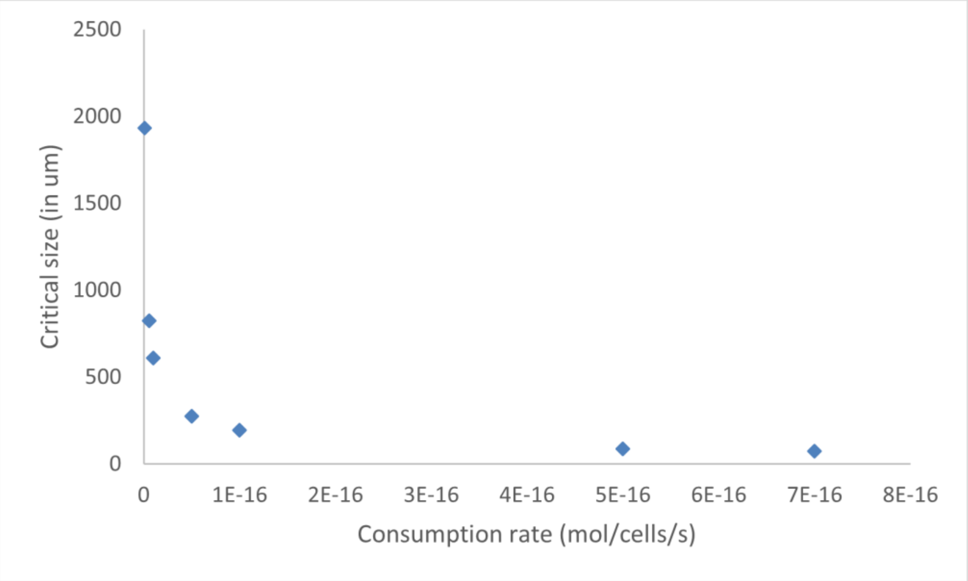


Our results may explain why 3D organoids exhibit premature cell death when they grow beyond 1 mm under a pure diffusive culturing condition. It is important to note that such critical radius is unique in 3D geometry and exists because of the establishment of the depleted boundary layer around the 3D organoid. 2D cell culture on a plane does not encounter such a transport limit.

### 5. Analysis of organoid growth under continuous microfluidic perfusion culturing by an intra-organoid diffusion-reaction model

To culture the organoid beyond the critical radius, a cross-chamber microfluidic perfusion strategy is proposed which aims to prevent the establishment of the depleted nutrient boundary layer and maintain the nutrient concentration around the organoid. In this section, we sought to understand the "healthy" growth dynamic of the organoid under the microfluidic perfusion condition. **By investigating an intra-organoid diffusion-reaction transport model, we show that this problem can be reduced to the analysis of a non-linear one-dimensional system:** $\dot{a} = f(a)$, where $\dot{a}$ is the first derivative of organoid radius $a$ with respect to time. **Using dimensional analysis, we demonstrate that under continuous microfluidic perfusion, the growth of the organoid follows a simple square root scaling of the culturing time.**

### 5.1. The growth of an organoid as a one-dimensional dynamic system

The growth of an organoid can be considered as transforming nutrients into new cells. The new cells grow from the existing organoid and hence increase the size of the organoid. *Note that a hidden assumption of the "healthy" organoid is that no cell death inside the organoid.* Translating to our model analysis, this assumption means that a cell, once grown, keeps the same volume and a consistent nutrient consumption rate, $\gamma_{cell}$. We also introduce a new parameter, $\eta_{cell}$, which denotes the total amount of nutrient required to grow a new cell (a nutrient-consumed unit volume). $\eta_{cell}$ has a unit of **mol (per cell)**.

We first assume a separation of timescales to set up the question. Two relevant timescales are discussed: the timescale for nutrient molecules to diffuse into the organoid center from outside for nutrient supply, $t_*$, and the timescale for a new cell to grow in the outer shell of the existing organoid, $t_{growth}$. From the experimental observation, the cell growth timescale $t_{growth}$ (on the order of days to weeks) is much larger than nutrient molecule diffusion timescale $t_*$ (on the order of hours). Therefore, we can separate these two timescales and consider the intra-organoid nutrient diffusion as a pseudo-steady-state process.

We then consider the intra-organoid nutrient mass balance within a small time interval. The diffusion-reaction transport process can be described as:

$$D_{organoid}\nabla^2 C^{in} = \gamma_{cell}\rho_{cell} + \eta_{cell}\rho_{cell}\frac{4\pi a^2 \dot{a}}{\frac{4}{3}\pi a^3} \quad \text{(S7)}$$

The LHS of the equation describes the diffusion-dominant transport of the critical nutrient inside the organoid, where $C^{in}$ denotes the intra-organoid concentration profile. The RHS of the equation describes the nutrient consumption of the organoid within this time interval. The first term, $\gamma_{cell}\rho_{cell}$, denotes the volume averaged nutrient consumption rate of existing cells in the organoid. The second term, $\eta_{cell}\rho_{cell}\frac{4\pi a^2 \dot{a}}{\frac{4}{3}\pi a^3}$, denotes the volume averaged nutrient consumption rate to grow new cells in the derivative outer shell of the organoid within this time interval. Under the pseudo-steady-state assumption, the second RHS term is a small perturbation compared to the first RHS term. Note that $\dot{\boldsymbol{a}}$ represents the derivative thickness of the newly grown outer shell of the organoid within this time interval. $\dot{\boldsymbol{a}} = \frac{da}{dt}$ has a unit of **m / s** and unifies the unit of all three terms in the equation.

To solve the intra-organoid concentration profile, we reorganize the equation:

$$\nabla^2 C^{in} = \Phi \quad \text{(S8)}$$

$$\Phi = \frac{\gamma_{cell}\rho_{cell} + 3\eta_{cell}\rho_{cell}\,\dot{\alpha}/\alpha}{D_{organoid}} \quad \text{(S9)}$$

We then specify the boundary conditions. Due to the continuous microfluidic perfusion culturing, the outside boundary condition at the outer edge of the organoid is expected to be at the fresh media concentration, $C_0$. The inside boundary condition at the center of the organoid can be specified by symmetry (Neumann boundary condition).

$$\begin{cases} C^{in}(r = a) = C_0 \\ \frac{dC^{in}}{dr}(r = 0) = 0 \end{cases} \quad (S10)$$

We can hence obtain the intra-organoid nutrient distribution:

$$C^{in}(r) = \frac{\Phi}{6}(r^2 - a^2) + C_0 \quad \text{(S11)}$$

We are interested in understanding the growth dynamic of the organoid, which can be analyzed by solving for $\dot{\boldsymbol{a}}$. To obtain the expression of $\dot{\boldsymbol{a}}$, we integrate nutrient concentration profile over the volume of the organoid for the total amount of the nutrient in the organoid, and assume it equals to a certain constant, $\boldsymbol{M}$. $\boldsymbol{M}$ has a unit of **mol**.

$$M = \iiint_V C^{in}\,dV = \int_0^{\pi}\int_0^{2\pi}\int_0^{a} C^{in}(r) r^2 \sin\varphi\, dr d\theta d\varphi = \frac{4\pi}{3} C_0 a^3 - \frac{4\pi}{45}\Phi a^5 \quad \text{(S12)}$$

Intuitively, $\boldsymbol{M}$ is determined by the nutrient supply and the scale of the organoid. For now, we don't know the value of $\boldsymbol{M}$ and we call it as the "nutrient supply constant". In later analysis, we will show that under the continuous microfluidic perfusion culturing, $\boldsymbol{M}$ does not affect the scaling of the growth dynamic.

By expanding $\boldsymbol{\Phi}$, we can easily solve for $\dot{\boldsymbol{a}}$ to obtain the dynamic relationship of this system:

$$\dot{a} = f(a) = \frac{\lambda_2}{\lambda_1} a^{-1} - \frac{\lambda_3}{\lambda_1} a - \frac{M}{\lambda_1} a^{-4} \quad \text{(S13)}$$

$$\lambda_1 = \frac{4\pi}{15 D_{organoid}} \eta_{cell}\rho_{cell};\ \lambda_2 = \frac{4\pi}{3} C_0;\ \lambda_3 = \frac{4\pi}{45 D_{organoid}} \gamma_{cell}\rho_{cell}. \quad \text{(S14)}$$

**5.2. Approximation of the organoid growth dynamic by dimensional analysis**

$\boldsymbol{f(a)}$ appears to be highly non-linear. Hence, we perform non-dimensionalization of the system to simplify the dynamic relationship.

We first make an estimation of nutrient supply constant $\boldsymbol{M}$. The nutrient concentration in the media and right outside of the organoid is held constant as $\boldsymbol{C_0}$ by the continuous microfluidic perfusion culturing condition. $\boldsymbol{C_0}$ is much larger than the consumption density of the organoid to ensure a sufficient nutrient condition and the nutrient concentration inside the organoid is always smaller than $\boldsymbol{C_0}$. Therefore, we can estimate the expression of $\boldsymbol{M}$ as: $\boldsymbol{M = \varepsilon C_0 V_{organoid}}$, where $\boldsymbol{\varepsilon}$ is a smaller-than-one number. The upper limit of $\boldsymbol{\varepsilon}$ can be estimated by the ratio between the nutrient consumption density and the nutrient supply density. Hence, we estimate $\boldsymbol{\varepsilon}$ as: $\boldsymbol{\varepsilon = \frac{\gamma_{cell}\rho_{cell}}{C_0 Q_{flow\,rate}/V_{trap}} = \frac{\gamma_{cell}\rho_{cell}t_{flow}}{C_0}}$. Here, we introduce a new time scale, $\boldsymbol{t_{flow} = \frac{V_{trap}}{Q_{flow\,rate}}}$, which denotes the characteristic timescale to refresh the media in a single microfluidic trap. We can qualitatively compare the three timescales involved in this process. $\boldsymbol{t_{flow}}$ is orders of magnitude smaller than $\boldsymbol{t_*}$ under the continuous microfluidic perfusion condition to allow for a sufficient nutrient delivery. $\boldsymbol{t_*}$ is further orders of magnitude smaller than $\boldsymbol{t_{growth}}$ as suggested by experimental evidence.

To perform non-dimensionalization, we denote $\boldsymbol{a_*}$ as a characteristic length scale of the organoid radius and choose $\boldsymbol{t_*}$ as the characteristic time scale. We especially select $\boldsymbol{a_*}$ so that $\boldsymbol{(a_*)^2 = D_{organoid}t_*}$. We can then render the dynamic equation $\boldsymbol{\dot{a} = f(a)}$ dimensionless by defining the non-dimensional organoid radius: $\boldsymbol{\bar{a} = a/a_*}$, and the non-dimensional time $\boldsymbol{\bar{t} = t/t_*}$. Substituting the estimated expression of $\boldsymbol{M}$ in the dynamic equation $\boldsymbol{\dot{a} = f(a)}$, we then obtain the dimensionless dynamic equation:

$$\boldsymbol{\frac{d\bar{a}}{d\bar{t}} = \frac{5C_0}{\eta_{cell}\rho_{cell}}(1-\varepsilon)\bar{a}^{-1} - \frac{1}{3}\frac{\gamma_{cell}}{\eta_{cell}}t_*\bar{a}} \quad \text{(S15)}$$

Since $\boldsymbol{\eta_{cell}}$ is the total amount of the nutrient required to grow a new cell and $\boldsymbol{\gamma_{cell}}$ is the single cell nutrient consumption rate, it is reasonable to suggest that $\boldsymbol{\eta_{cell}/\gamma_{cell}}$ is at the same order of $\boldsymbol{t_{growth}}$. Due to the abundant nutrient in the media, the density of nutrient required to grow a new cell $\boldsymbol{\eta_{cell}\rho_{cell}}$ is at the same order of the nutrient supply concentration $\boldsymbol{C_0}$. Hence, the coefficient of the second RHS term $\boldsymbol{\frac{1}{3}\frac{\gamma_{cell}}{\eta_{cell}}t_*}$ is at the same order of $\boldsymbol{\frac{1}{3}\frac{t_*}{t_{growth}}}$, while orders of magnitude smaller than the coefficient of the first RHS term $\boldsymbol{\frac{5C_0}{\eta_{cell}\rho_{cell}}(1-\varepsilon)}$. We approximate the dimensionless dynamic equation as:

$$\frac{d\bar{a}}{d\bar{t}} \approx \frac{5C_0}{\eta_{cell}\rho_{cell}}(1-\varepsilon)\bar{a}^{-1} = \frac{5C_0}{\eta_{cell}\rho_{cell}}\left(1-\frac{\gamma_{cell}\rho_{cell}t_{flow}}{C_0}\right)\bar{a}^{-1} \quad \text{(S16)}$$

Therefore, we can solve the approximated dynamic equation to obtain the scaling relationship between the radius of the organoid and the culturing time:

$$a \approx \sqrt{\frac{10D_{organoid}C_0}{\eta_{cell}\rho_{cell}}\left(1-\frac{\gamma_{cell}\rho_{cell}t_{flow}}{C_0}\right)t} \quad \text{(S17)}$$

For organoids cultured with sufficient nutrient delivery under the continuous microfluidic perfusion condition, the growth dynamic should follow such a scaling relationship. It hence provides us a guideline to analyze the experimental observation of the organoid. We can apply natural log to both sides of the equation, and obtain:

$$\ln a \approx \frac{1}{2}\ln t + \frac{1}{2}\ln\frac{10D_{organoid}C_0}{\eta_{cell}\rho_{cell}} + \frac{1}{2}\ln\left(1-\frac{\gamma_{cell}\rho_{cell}t_{flow}}{C_0}\right) \quad \text{(S18)}$$

If we apply log to experimental data and plot $\ln a$ versus $\ln t$, **the well-fed** _(no outside nutrient depletion layer)_ **and healthy** _(no premature cell death)_ **organoids should show a linear $\ln a - \ln t$ relationship with a ½ slope**. The experimental data shows **non-linear relationship or non-½ slope indicates that the organoid was nutrient-deficient or underwent significant intra-organoid cell death at some point during the culturing**. The intercept of the $\ln a - \ln t$ relationship shown in the second RHS term may indicate certain physiological properties of the organoid. For example, $D_{organoid}$ may indicate the integrity of the organoid which is related to how dense the ECM is and hence the intra-organoid diffusivity; $\eta_{cell}$ and $\rho_{cell}$ may indicate the cell type and the differentiation direction of the organoid. These physiological properties are expected to be affected by many factors, including the overall nutrient delivery, ratio between different nutrients, shear stress during culturing, etc. This is not captured in this model but can be experimentally determined. The intercept of the $\ln a - \ln t$ relationship shown in the third RHS term is affected by the culturing condition, namely the flow rate of the microfluidic perfusion. However, upon close inspection, we find that **as long as the perfusion ensures an abundant nutrient environment without the depletion boundary layer, the influence on the organoid growth dynamic by the flow rate is almost negligible**. This is because $\frac{\gamma_{cell}\rho_{cell}}{C_0}$ is at the same order of magnitude of $t_*$. The third RHS is close $\frac{1}{2}\ln\left(1-\frac{t_{flow}}{t_*}\right)$, and hence $\approx \frac{1}{2}\ln(1) = 0$.

**Figure S3. Convective-based culturing strategy enables sufficient nutrient delivery and robust organoid culture** ((a. Top: Schematic describing culture set-up for syringe-based recirculation of media using convection-based device. The inlet of the device is connected to a syringe pump for perfusion. The outlet of the device is connected to a second syringe barrel which serves as a reservoir and enables the mixing of spent media with fresh media from the pump. Created with Biorender.com. Two different flow conditions were tested. Multi-step recycle; where the forward and reverse flow rates are uneven and single-step recycle; where the forward and reverse flow rates are even. The total volume dispensed in both forward and reverse directions were kept constant for all flow conditions. Bottom:  Log-log plot of all convection-based culturing methods highlighting the convergence of the convection-based methods around 0.5 b. Box-whisker plot of various brightfield metrics characterizing organoid quality in multi-step media recycle at different flow rates (diameter, solidity, circularity, and aspect ratio). n = 13 organoids for 1ml $hr^{-1}$ condition. n = 20 organoids for 2ml $hr^{-1}$ condition. Using a two-tailed unpaired t-test with Welch-correction for the 2 groups. p value = 0.7405, 0.2085, 0.2851, 0.1421 for diameter, circularity, aspect ratio and solidity respectively))

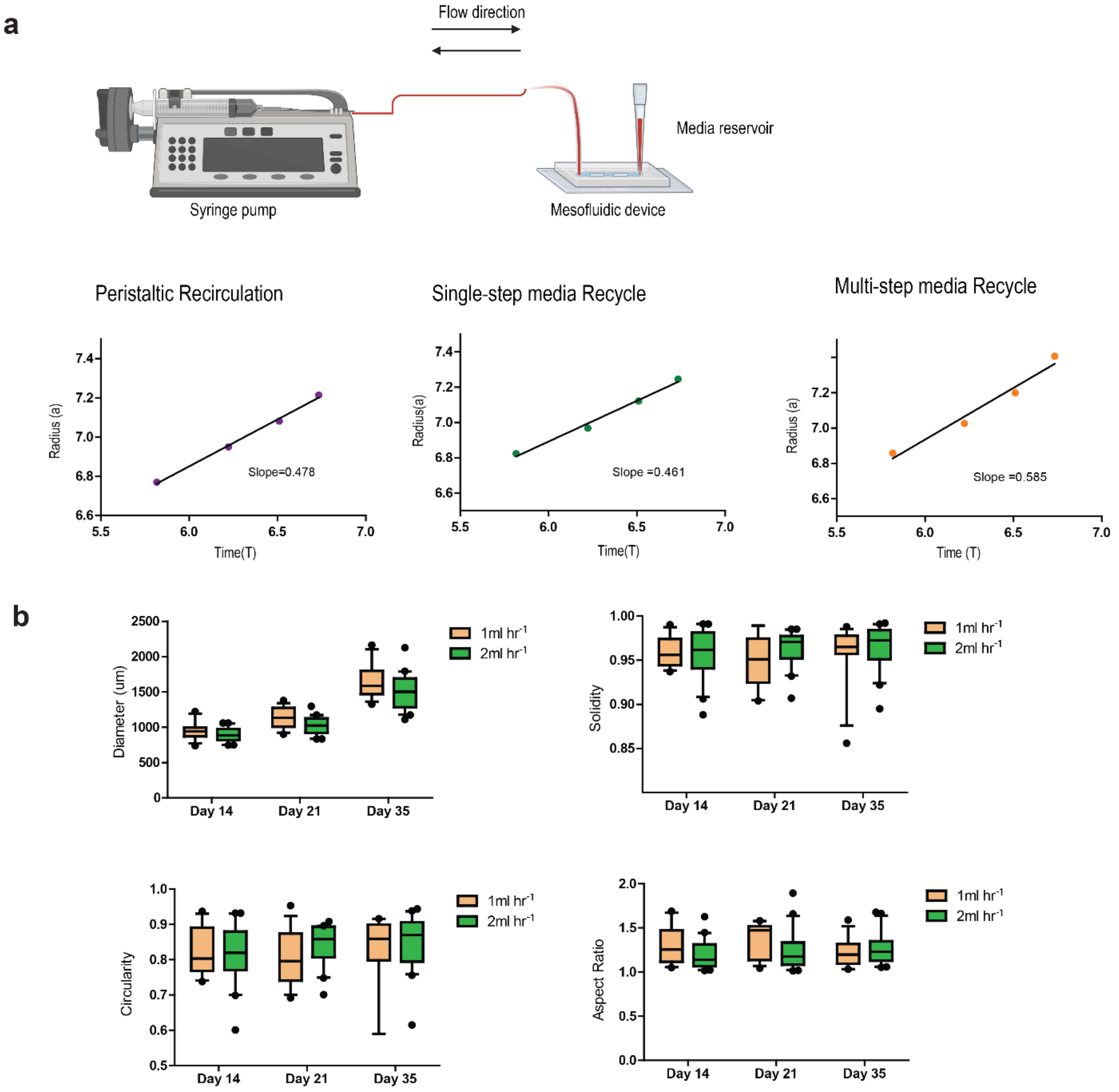
a
Flow direction
Media reservoir
Syringe pump
Mesofluidic device
Peristaltic Recirculation
Single-step media Recycle
Multi-step media Recycle
Radius (a)
Time(T)
Slope=0.478
Radius(a)
Slope =0.461
Slope =0.585
Time (T)
b
Diameter (um)
Solidity
Circularity
Aspect Ratio
Day 14
Day 21
Day 35
1ml hr-1
2ml hr-1

**Figure S4. Characterization of intra-device variability in the mesofluidic bioreactor under peristaltic perfusion (((**a). Organoid diameter as a function of position in the device. Diameters of day 14 and day 35 organoids from 7 devices from 3 independent experiments are shown. (b). Organoid circularity as a function of position in the device. Circularity values of day 14 and day 35 organoids from 7 devices from 3 independent experiments are shown. (c). Organoid solidity as a function of position in the device. Solidity values of day 14 and day 35 organoids from 7 devices from 3 independent experiments are shown. (d). Organoid aspect ratio as a function of position in the device. Aspect ratio values of day 14 and day 35 organoids from 7 devices from 3 independent experiments are shown. Each device is indicated by a different symbol and color. Each independent experiment comprises of 2-3 devices containing 6-8 organoids.))

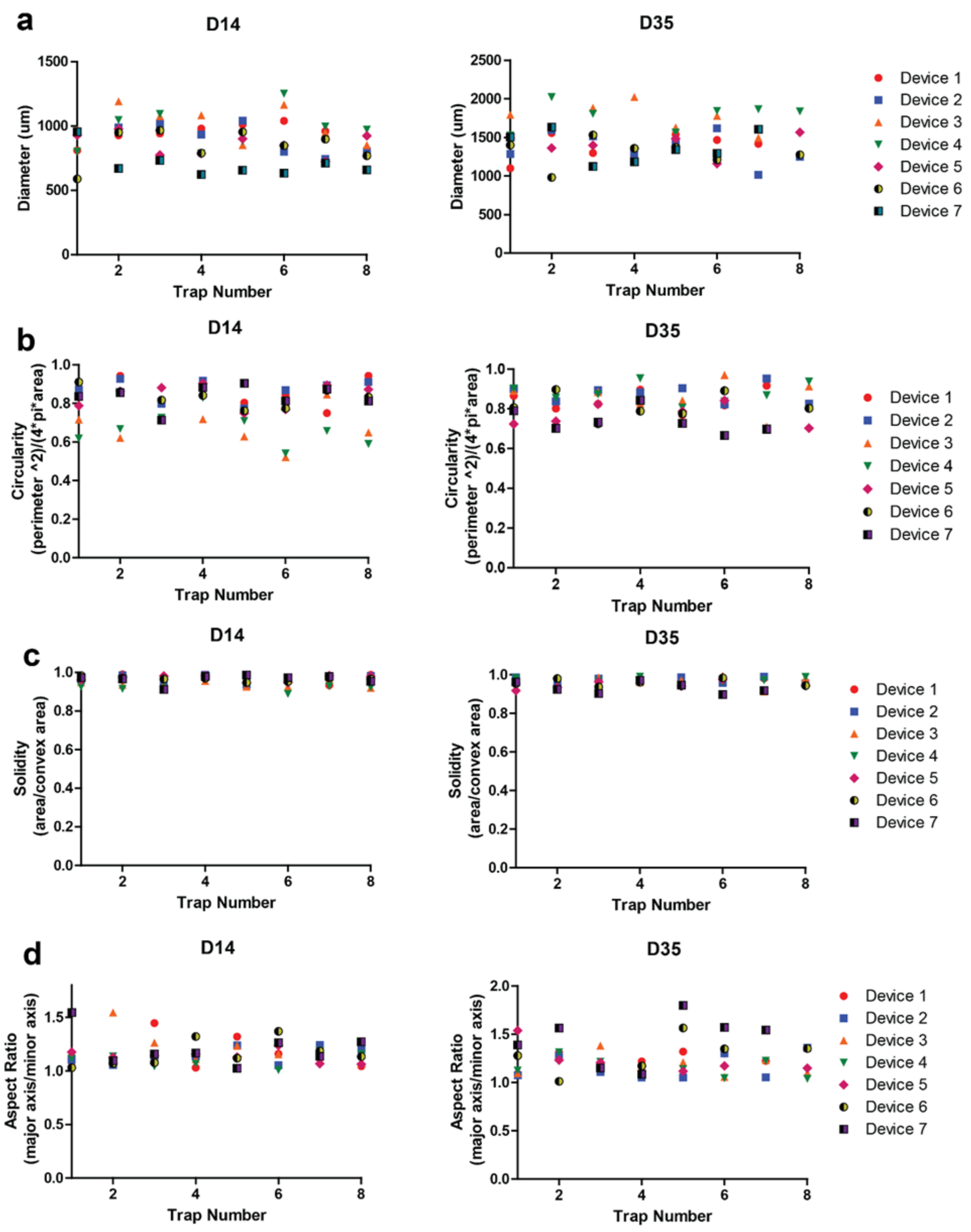
a
D14
D35
Diameter (um)
Trap Number
Device 1
Device 2
Device 3
Device 4
Device 5
Device 6
Device 7
b
Circularity
(perimeter ^2)/(4*pi*area)
c
Solidity
(area/convex area)
d
Aspect Ratio
(major axis/minor axis)

**Figure S5. Characterization of intra-device variability in the mesofluidic bioreactor under syringe perfusion (multi-step recycle)** a. Organoid diameter as a function of position in the device. Diameters of day 14 and day 35 organoids from 5 devices from 2 independent experiments are shown. b. Organoid circularity as a function of position in the device. Circularity values of day 14 and day 35 organoids from 5 devices from 2 independent experiments are shown. c. Organoid solidity as a function of position in the device. Solidity values of day 14 and day 35 organoids from 5 devices from 2 independent experiments are shown. d. Organoid aspect ratio as a function of position in the device. Aspect ratio values of day 14 and day 35 organoids from 5 devices from 2 independent experiments are shown. Each device is indicated by a different symbol and color. Each independent experiment comprises of 2-3 devices containing 6-8 organoids.

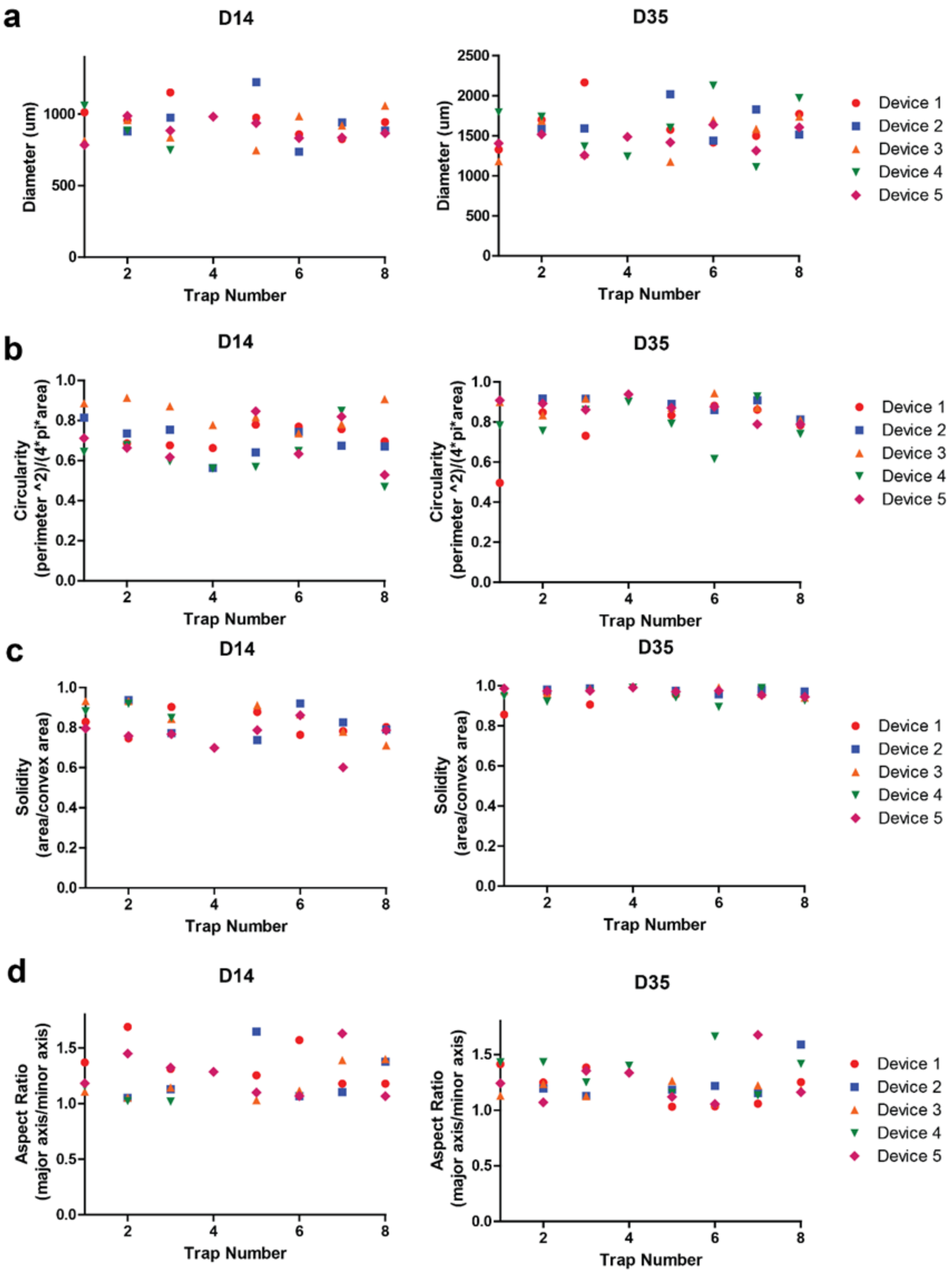
a
D14
D35
Diameter (um)
Trap Number
Device 1
Device 2
Device 3
Device 4
Device 5
b
Circularity
(perimeter ^2)/(4*pi*area)
c
Solidity
(area/convex area)
d
Aspect Ratio
(major axis/minor axis)

**Table S2. Summary of features obtained from bright field images which used in this study.**

| No. | Feature Name |
|---|---|
| 1 | Individual Scaling Factors |
| 2 | Deviation_from_scaling_absolute |
| 3 | Deviation_from_scaling_normalized |
| 4 | d14_Area (um2) |
| 5 | d14_Diameter (um) |
| 6 | d14_circularity |
| 7 | d14_perimeter (um) |
| 8 | d14_aspect ratio |
| 9 | d14_solidity |
| 10 | d14_intensity_mean |
| 11 | d14_intensity_std |
| 12 | d21_Area (um2) |
| 13 | d21_Diameter (um) |
| 14 | d21_circularity |
| 15 | d21_perimeter (um) |
| 16 | d21_aspect ratio |
| 17 | d21_solidity |
| 18 | d21_intensity_mean |
| 19 | d21_intensity_std |
| 20 | d28_Area (um2) |
| 21 | d28_Diameter (um) |
| 22 | d28_circularity |
| 23 | d28_perimeter (um) |
| 24 | d28_aspect ratio |
| 25 | d28_solidity |
| 26 | d28_intensity_mean |
| 27 | d28_intensity_std |
| 28 | d35_Area (um2) |
| 29 | d35_Diameter (um) |
| 30 | d35_circularity |
| 31 | d35_perimeter (um) |
| 32 | d35_aspect ratio |
| 33 | d35_solidity |
| 34 | d35_intensity_mean |
| 35 | d35_intensity_std |

**Figure S6. High-content brightfield images of organoid in culture can be used to determine the biological quality of organoids** ((a. Plot showing consensus data for manual curation. b. Plot showing differences in all features tested between organoids that develop normally and defective organoids. Data is representative of 15 devices containing 6-8 organoids obtained from all the perfusion conditions tested with the platform ))

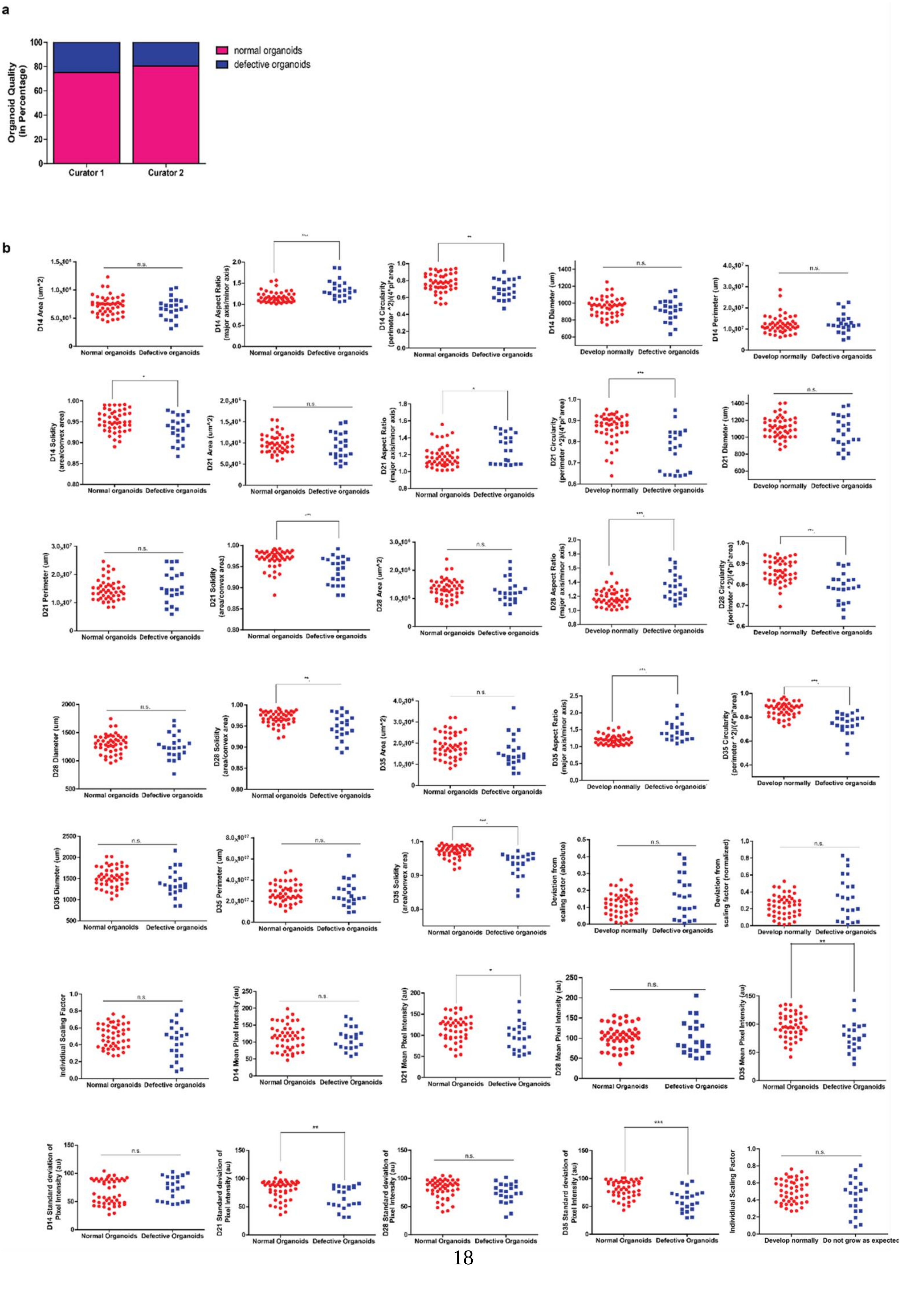
a
Organoid Quality (in Percentage)
normal organoids
defective organoids
Curator 1
Curator 2
b
D14 Area (um^2)
D14 Aspect Ratio (major axis/minor axis)
D14 Circularity (perimeter ^2)/(4*pi*area)
D14 Diameter (um)
D14 Perimeter (um)
D14 Solidity (area/convex area)
D21 Area (um^2)
D21 Aspect Ratio (major axis/minor axis)
D21 Circularity (perimeter ^2)/(4*pi*area)
D21 Diameter (um)
D21 Perimeter (um)
D21 Solidity (area/convex area)
D28 Area (um^2)
D28 Aspect Ratio (major axis/minor axis)
D28 Circularity (perimeter ^2)/(4*pi*area)
D28 Diameter (um)
D28 Solidity (area/convex area)
D35 Area (um^2)
D35 Aspect Ratio (major axis/minor axis)
D35 Circularity (perimeter ^2)/(4*pi*area)
D35 Diameter (um)
D35 Perimeter (um)
D35 Solidity (area/convex area)
Deviation from scaling factor (absolute)
Deviation from scaling factor (normalized)
Individual Scaling Factor
D14 Mean Pixel Intensity (au)
D21 Mean Pixel Intensity (au)
D28 Mean Pixel Intensity (au)
D35 Mean Pixel Intensity (au)
D14 Standard deviation of Pixel Intensity (au)
D21 Standard deviation of Pixel Intensity (au)
D28 Standard deviation of Pixel Intensity (au)
D35 Standard deviation of Pixel Intensity (au)
Normal organoids
Defective organoids
Develop normally
Do not grow as expected
n.s.

**Figure S7. High-content brightfield images of organoid in culture can be used to determine the biological quality of organoids** ((3D rendering of organoids cultured in the microfluidic platform showing distribution of rosette structures and surrounding neuronal marker (TUJ1)).

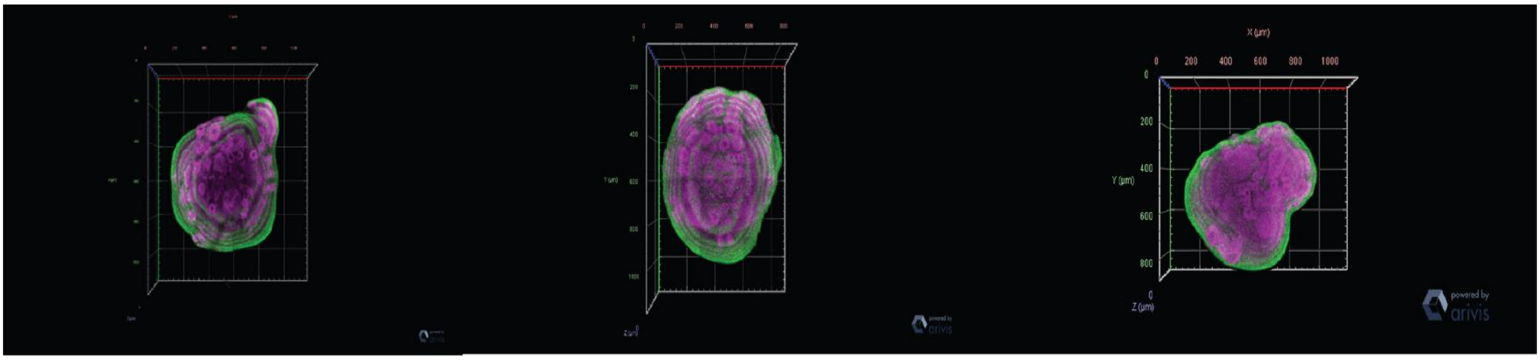

**Figure S8. The molecular signature of the pre-selected organoids cultured in the mesofluidic platform shows normal maturation of cell phenotypes in 3D organoids** ((Quantification of organoid rosette number for day 28 and day 35 organoids cultured under convective-based culture method vs. spin omega. Using a 2-way ANOVA with Bonferroni correction. where ** indicates p-value <0.01. Data is representative of 6-8 organoids and 2 devices per experiment group))

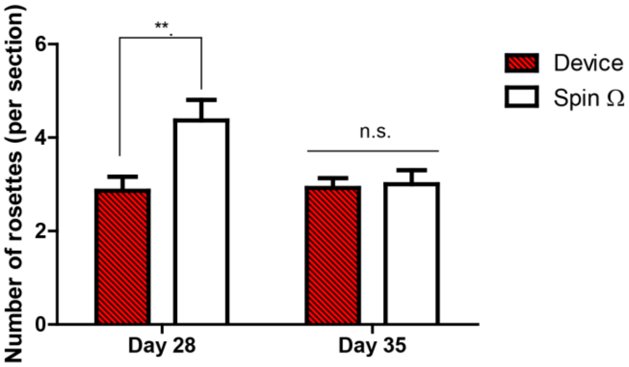

**Figure S9. Characterization of inter-device variability in the mesofluidic bioreactor under peristaltic perfusion**. (((a). Box-Whisker plot showing the variance in number of neural rosettes in day 28 organoid slices across two devices. n = 13 slices for device 1. n = 16 slices for device 2. Using an F-test to compare variances, p-value= 0.1747. (b) Box-Whisker plot showing the variance in number of neural rosettes in organoid slices across two devices for day 35 organoids. n =18 slices for device 1. n= 9 slices for device 2. n = 21 slices for device 3. Using a one-way ANOVA (with Kruskal-Wallis correction) analysis, p-value = 0.3387. (c). Box-Whisker plot showing the variance in number of Sox2+ cells in day 28 organoid slices across two devices. n = 13 slices for device 1. n = 25 slices for device 2. Using an F-test to compare variances, p-value= 0.4606. (d) Box-Whisker plot showing the variance in number of TBR2+ cells in day 28 organoid slices across two devices. n =4 slices for device 1. n = 8 slices for device 2. Using an F-test to compare variances, p-value=0.6249. (e) Box-Whisker plot showing the variance in number of CTIP2+ cells in day 28 organoid slices across two devices. n = 4 slices for device 1. n = 8 slices for device 2. Using an F-test to compare variances, p-value=0.0615.))

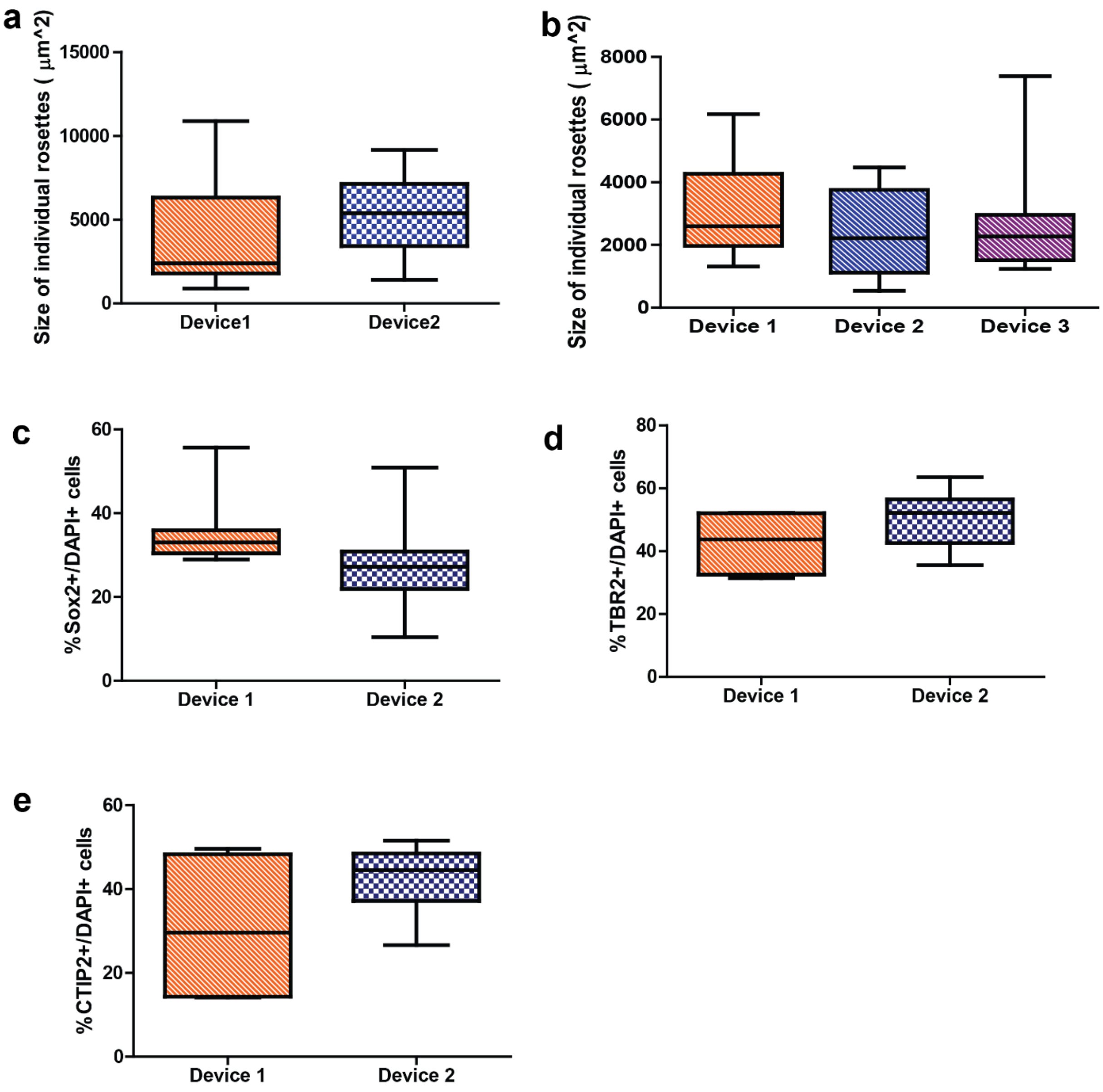
a
Size of individual rosettes ( µm^2)
15000
10000
5000
0
Device1
Device2
b
Size of individual rosettes ( µm^2)
8000
6000
4000
2000
0
Device 1
Device 2
Device 3
c
%Sox2+/DAPI+ cells
60
40
20
0
Device 1
Device 2
d
%TBR2+/DAPI+ cells
80
60
40
20
0
Device 1
Device 2
e
%CTIP2+/DAPI+ cells
60
40
20
0
Device 1
Device 2

**Figure S10. Correlation of global transcriptomes between human forebrain organoids and fetal human brain development** ((Shown are heatmaps of Pearson's correlation analysis of RNA-seq datasets among human forebrain organoids at different stages cultured in the mesofluidic and SpinΩ platforms and published transcriptome datasets of 3 different human cortical subregions, including ventrolateral prefrontal cortex (VLPFC; a), orbital frontal cortex (OFC; b), and dorsolateral prefrontal cortex (DLPFC; c), at 4 developmental stages from Allen Brain Atlas. Data are from 2 devices per experimental group and 4-5 organoids per device.))

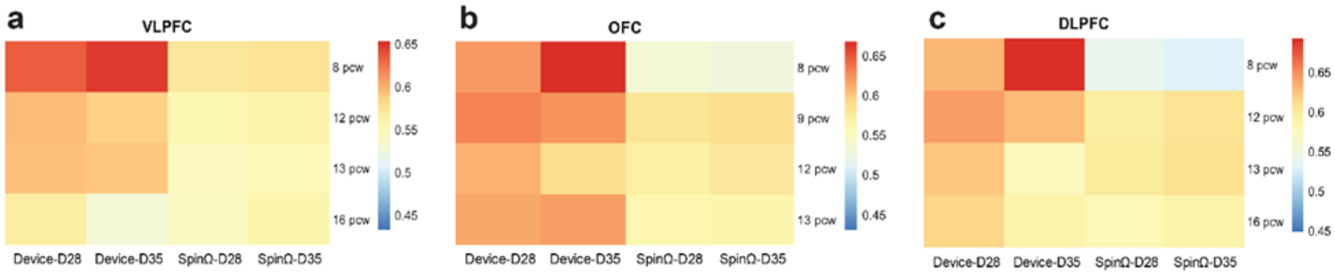

**Table S3. Summary of hyper-parameters tested in cross-validation/grid-search process for the SVM model.**

| Kernel | Gamma | C | Degree | Mean 5-fold CV score (f1) |
|---|---|---|---|---|
| rbf | 0.1 | 1 | - | 0.812428 |
| rbf | 0.01 | 1 | - | 0.91128 |
| rbf | 0.001 | 1 | - | 0.809144 |
| rbf | 0.0001 | 1 | - | 0.809144 |
| rbf | 0.1 | 10 | - | 0.814441 |
| rbf | 0.01 | 10 | - | 0.902772 |
| rbf | 0.001 | 10 | - | 0.91287 |
| rbf | 0.0001 | 10 | - | 0.809155 |
| rbf | 0.1 | 100 | - | 0.814441 |
| rbf | 0.01 | 100 | - | 0.896537 |
| rbf | 0.001 | 100 | - | 0.900625 |
| rbf | 0.0001 | 100 | - | 0.909515 |
| rbf | 0.1 | 1000 | - | 0.814441 |
| rbf | 0.01 | 1000 | - | 0.896537 |
| rbf | 0.001 | 1000 | - | 0.888208 |
| rbf | 0.0001 | 1000 | - | 0.895197 |
| linear | - | 1 | - | 0.882952 |
| linear | - | 10 | - | 0.882939 |
| linear | - | 100 | - | 0.882939 |
| linear | - | 1000 | - | 0.882939 |
| poly | 0.1 | 1 | 2 | 0.819891 |
| poly | 0.01 | 1 | 2 | 0.809115 |
| poly | 0.001 | 1 | 2 | 0.809144 |
| poly | 0.0001 | 1 | 2 | 0.809144 |
| poly | 0.1 | 1 | 3 | 0.864415 |
| poly | 0.01 | 1 | 3 | 0.809144 |
| poly | 0.001 | 1 | 3 | 0.809144 |
| poly | 0.0001 | 1 | 3 | 0.809144 |
| poly | 0.1 | 10 | 2 | 0.814939 |
| poly | 0.01 | 10 | 2 | 0.833966 |
| poly | 0.001 | 10 | 2 | 0.809144 |
| poly | 0.0001 | 10 | 2 | 0.809144 |
| poly | 0.1 | 10 | 3 | 0.864463 |
| poly | 0.01 | 10 | 3 | 0.844481 |
| poly | 0.001 | 10 | 3 | 0.809144 |
| poly | 0.0001 | 10 | 3 | 0.809144 |
| poly | 0.1 | 100 | 2 | 0.814939 |
| poly | 0.01 | 100 | 2 | 0.819891 |
| poly | 0.001 | 100 | 2 | 0.809115 |
| poly | 0.0001 | 100 | 2 | 0.809144 |
| poly | 0.1 | 100 | 3 | 0.864463 |
| poly | 0.01 | 100 | 3 | 0.862356 |
| poly | 0.001 | 100 | 3 | 0.809144 |

| | | | | |
|---|---|---|---|---|
| poly | 0.0001 | 100 | 3 | 0.809144 |
| poly | 0.1 | 1000 | 2 | 0.814939 |
| poly | 0.01 | 1000 | 2 | 0.814939 |
| poly | 0.001 | 1000 | 2 | 0.833966 |
| poly | 0.0001 | 1000 | 2 | 0.809144 |
| poly | 0.1 | 1000 | 3 | 0.864463 |
| poly | 0.01 | 1000 | 3 | 0.864415 |
| poly | 0.001 | 1000 | 3 | 0.809144 |
| poly | 0.0001 | 1000 | 3 | 0.809144 |

**Figure S11. Figure showing SVM performance on both the training and test data sets as the size of the training data set is increased.** ((The shaded area represents the standard deviation from the mean F1 score, which helps us understand the variability in the performance measurements. From the graph, we observed that the F1 score on the test set (the 21 images we kept separate for testing) reached a stable value once the training set size reached 40. This stable score indicated that the SVM model was not overfitting when trained with more than 40 images, meaning it was generalizing well to new, unseen data))

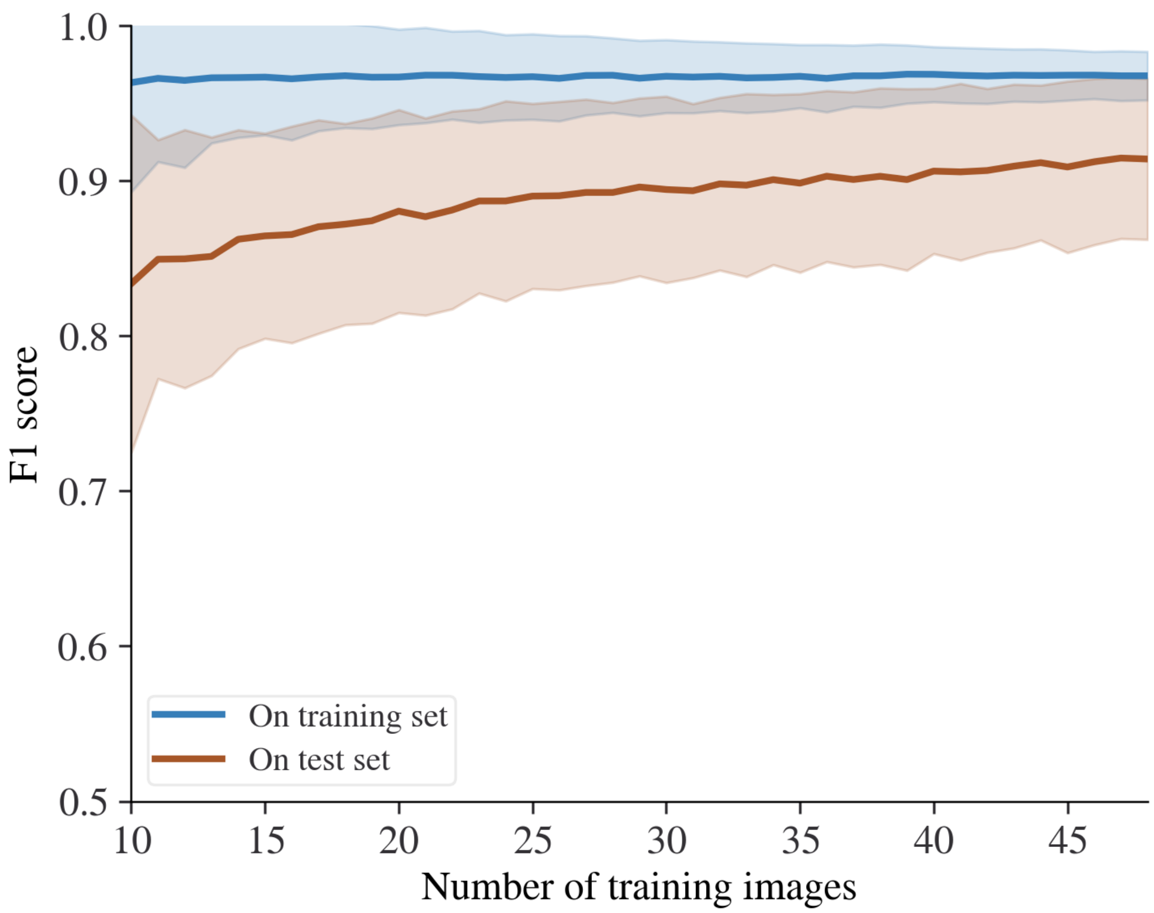

**References**


[1] McMurtrey, R. J., *Tissue engineering. Part C, Methods* **2016,** *22* (3), 221-249. DOI 10.1089/ten.TEC.2015.0375.
[2] Greenspan, H. P., *Studies in Applied Mathematics* **1972,** *51* (4), 317-340. DOI https://doi.org/10.1002/sapm1972514317.
[3] Maggelakis, S. A.; Adam, J. A., *Mathematical and Computer Modelling* **1990,** *13* (5), 23-38. DOI 10.1016/0895-7177(90)90040-T.
[4] Maggelakis, S. A., *Mathematical and Computer Modelling* **1993,** *17* (10), 19-29. DOI 10.1016/0895-7177(93)90114-E.
[5] Varum, S.; Rodrigues, A. S.; Moura, M. B.; Momcilovic, O.; Easley 4th, C. A.; Ramalho-Santos, J.; Van Houten, B.; Schatten, G., *PloS one* **2011,** *6* (6), e20914-e20914. DOI 10.1371/journal.pone.0020914.
[6] Herculano-Houzel, S., *PLOS ONE* **2011,** *6* (3), e17514.
[7] Karbowski, J., *BMC Biology* **2007,** *5* (1), 18-18. DOI 10.1186/1741-7007-5-18.
[8] Berger, E.; Magliaro, C.; Paczia, N.; Monzel, A. S.; Antony, P.; Linster, C. L.; Bolognin, S.; Ahluwalia, A.; Schwamborn, J. C., *Lab on a Chip* **2018,** *18* (20), 3172-3183. DOI 10.1039/C8LC00206A.
